# Spatiotemporal unfolding of prefrontal neural response divergence in adolescent depression during naturalistic experience

**DOI:** 10.64898/2026.06.04.730065

**Authors:** Qianhui Liu, Weiyang Shi, Haiyan Wang, Wenlu Li, Nianyi Liu, Siyao Dong, Congying Chu, Lingzhong Fan, Tianzi Jiang

## Abstract

Depression alters how emotional experience is interpreted and regulated, yet how depression-related neural divergence unfolds during ongoing experience remains poorly understood, leaving unresolved whether such divergence reflects stable trait-like abnormalities or state-dependent responses that emerge only at particular moments. This question is especially consequential in adolescence, when depression often emerges amid heightened social sensitivity and continuing maturation of prefrontal regulatory systems. Here, we used naturalistic film watching fMRI data of 36 adolescents with depression and 36 matched controls to map where, how, and when neural responses diverge between diagnostic groups. Neural polarization analysis localized group-level divergence to two prefrontal hubs, the dmPFC-rACC and left dlPFC, where individual similarity to the depression-group response pattern was associated with depressive symptom severity across the cohort. These local divergence sites were not spatially isolated. Seed-based inter-subject functional connectivity showed that they were embedded within distributed stimulus-locked coupling patterns spanning prefrontal control, salience, and subcortical reward circuits. Time-resolved phase synchrony further revealed that this divergence was not continuous, but arose at specific narrative moments, with the dmPFC-rACC diverging during socio-emotional events and the left dlPFC during cognitively demanding ones. Together, these findings move beyond static localization to reveal depression-related neural divergence as a spatially organized, network-embedded, and temporally gated response pattern shaped by the unfolding content of naturalistic experience.

## Introduction

Depression is characterized not only by persistent low mood, but also by systematic alterations in how emotionally meaningful experiences are interpreted, attended, and regulated. Depressed individuals preferentially elaborate negative information, struggle to disengage from emotionally aversive content, and show impaired use of positive information for mood regulation [1]. These alterations are especially consequential during adolescence, a developmental period marked by heightened emotional sensitivity and rapidly evolving social cognition, during which major depressive disorder becomes highly prevalent and strongly associated with self-harm and suicide [2-4]. Despite extensive neuroimaging research, how the depressed adolescent brain diverges from the healthy brain during continuous emotional experience remains poorly understood.

Neuroimaging studies have consistently linked depression to abnormal interactions between limbic systems involved in emotional salience and prefrontal systems supporting cognitive control [5-10]. During emotional tasks, depressed individuals show exaggerated responses to negative stimuli together with altered recruitment of insular, cingulate, striatal, and thalamic regions [6, 7]. Resting-state studies further demonstrate disrupted functional organization across the default-mode, salience, and frontoparietal control networks [8-10]. However, these approaches capture only partial aspects of emotional processing. Task paradigms typically isolate brief and decontextualized emotional operations, whereas resting-state paradigms examine intrinsic neural dynamics independent of external experience. By design, neither directly addresses how neural responses unfold during continuous, temporally evolving, emotionally and socially embedded experience.

Naturalistic paradigms, including movie watching and spoken narratives, enable the study of brain function under temporally continuous and emotionally contextualized conditions while retaining experimental control [11-13]. Because all participants are exposed to the same evolving stimulus, these paradigms permit direct quantification of inter-individual alignment in neural responses. Inter-subject correlation (ISC) and related approaches demonstrate that such alignment reflects not only shared sensory input, but also high-order psychological factors including interpretation, social affiliation, and affective traits [14-20]. Individuals sharing similar interpretations of ambiguous narratives exhibit convergent activity in high-order cortical systems, whereas trait-level characteristics such as paranoia selectively reshape responses to socially salient content [18-20]. Emerging evidence further indicates atypical neural alignment during naturalistic stimulation in depression. Adults with melancholic depression show reduced inter-subject synchrony during emotional film viewing [21], while adolescents with greater depressive symptom severity exhibit increasingly idiosyncratic responses to emotional narratives [22]. Altered recruitment of large-scale attention and control networks has likewise been observed in clinically depressed adolescents during movie watching [23].

Together, these findings suggest that neural responses in depression diverge from normative patterns during continuous experience. However, a central unresolved question is whether neural divergence in depression reflects a stable trait-like state or context-dependent responses that emerge only during particular moments of ongoing experience. This question is especially important in naturalistic settings, where emotional and cognitive demands continuously fluctuate across the narrative. Depression-related abnormalities may therefore not be continuously expressed, but instead emerge selectively during emotionally salient, socially charged, or cognitively demanding events.

Addressing this question requires approaches capable of resolving neural divergence at the timescale of ongoing experience. Existing inter-subject analyses can establish whether depressed and non-depressed individuals differ during naturalistic stimulation, but are less suited to determining when such divergence emerges across the evolving narrative. Neural polarization frameworks isolate variance associated with group-specific interpretation by quantifying the extent to which neural responses are more synchronized within groups than between groups [24, 25]. Extending these frameworks to the temporal domain is necessary, yet most dynamic implementations rely on sliding-window correlations that temporally smooth neural signals over tens of seconds, limiting sensitivity to transient narrative events [26]. Here, to overcome this limitation, we combined neural polarization with inter-subject phase synchrony (ISPS), which estimates moment-to-moment synchronization directly from the instantaneous phase of the BOLD signal [27]. This approach enables neural data themselves to identify the specific narrative contexts in which depression-related divergence emerges, rather than imposing predefined temporal boundaries or emotional categories.

Using this framework, we tested whether neural divergence in adolescent depression is spatially organized and selectively emerges during specific forms of ongoing experience (Fig. 1a). We combined neural polarization, inter-subject functional connectivity, and time-resolved inter-subject phase synchrony analyses of the fMRI data acquired while adolescents from the Healthy Brain Network [28] viewed an emotionally rich clip from Despicable Me (Fig. 1b). Two prefrontal regions, the dorsomedial prefrontal and rostral anterior cingulate cortex and the left dorsolateral prefrontal cortex, exhibited significant depression-related divergence and were associated with symptom severity across the cohort. Importantly, the two regions showed dissociable temporal profiles across the evolving narrative. Divergence in the dorsomedial prefrontal and rostral anterior cingulate cortex peaked during socio-emotional events, whereas divergence in the left dorsolateral prefrontal cortex emerged preferentially during cognitively demanding moments. Together, these findings reveal that neural divergence in adolescent depression is anatomically organized and dynamically shaped by narrative context, rather than continuously expressed throughout ongoing experience. By linking moment-to-moment neural divergence to specific forms of narrative context, this framework provides a naturalistic approach for studying how depression-related abnormalities unfold during continuous experience.

**Fig. 1.**
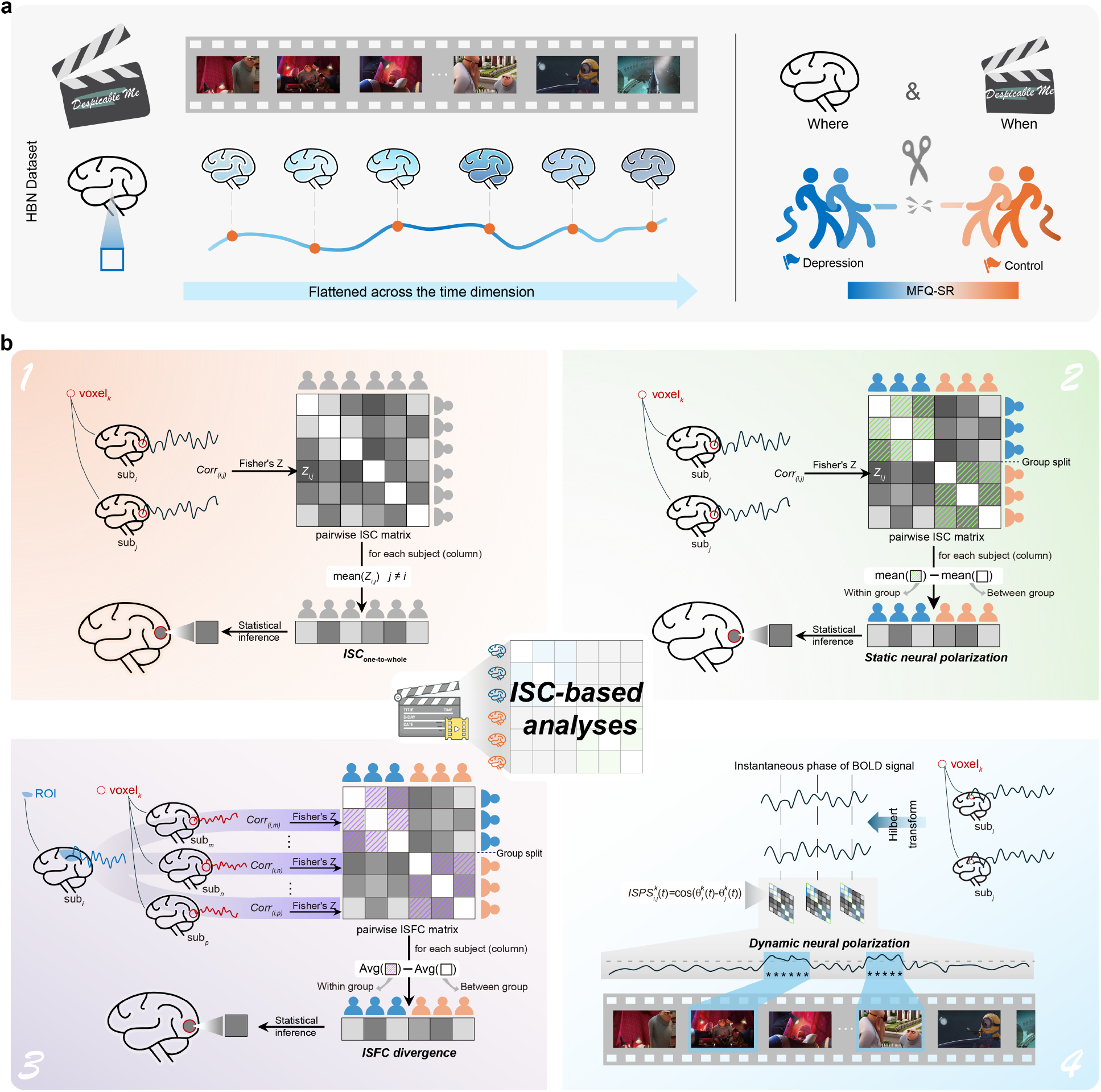
Study design and analytical framework. **a** Schematic of the experimental design. The analysis sample, drawn from the Healthy Brain Network (HBN) project, consisted of 36 participants with depression and 36 matched controls. Functional MRI data were acquired while participants viewed a 10-min clip from the movie *Despicable Me*. **b** Overview of the ISC-based analytical pipeline, comprising four progressive stages (voxel size enlarged for visualization). All analyses are built on a pairwise approach. (**1**) *Shared neural response analysis*. Each participant’s ISC value was computed as the average pairwise correlation with all other participants, regardless of group membership. (**2**) *Static neural polarization analysis*. Pairwise correlations were partitioned into within-group (green hatched blocks) and between-group components and the polarization index was defined as their difference. (**3**) *ISFC divergence analysis*. Voxel clusters identified in (2) served as seed regions. For each participant, the seed time course was correlated with whole-brain voxel-wise time courses of all other participants. The divergence index was computed as the difference between within-group (purple hatched blocks) an0064 between-group ISFC. (**4**) *Dynamic neural polarization analysis*. A time-resolved polarization was derived frame-wise by subtracting average between-group ISPS from average within-group ISPS. ISC, inter-subject correlation; ISFC, inter-subject functional connectivity; ISPS, inter-subject phase synchrony.

## Results

### Participants

The final analytical sample comprised 72 participants, including 36 individuals with a clinical diagnosis of depression and 36 matched controls. There were no significant differences between the two groups in terms of age, sex, psychiatric comorbidities or mean head motion during scanning (all *p* > 0.70; Table 1). Depression symptom severity, quantified using the self-report responses to the Moods and Feelings Questionnaire (MFQ-SR) [29], was significantly elevated in the depression group relative to controls (*t* (70)= 4.254, *p =* 6.40 × 10^−5^). Detailed demographic and clinical profiles are summarized in Table 1, with participant inclusion and matching protocols described in the *Methods*.

**Table 1.**
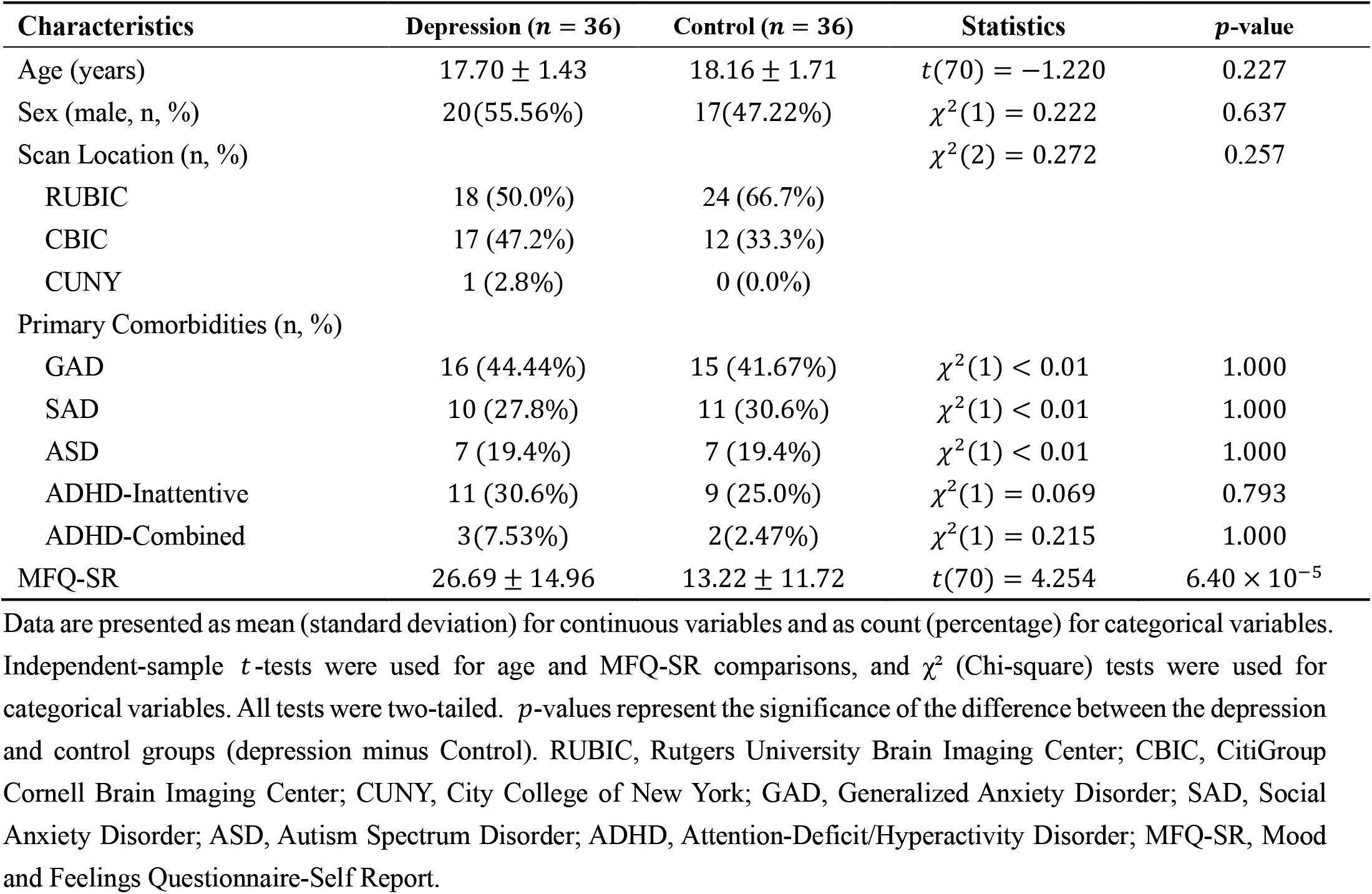
Demographic and clinical characteristics of depressed participants and matched controls.

### Emotional movie viewing elicits robust inter-subject neural synchrony

To verify that the naturalistic paradigm effectively engaged neural processing [14, 15, 17], we first assessed shared neural responses across the entire cohort (N=72). Regardless of diagnostic status, the emotional film clip elicited widespread neural synchrony primarily in unimodal sensory and downstream perceptual regions (Fig. 2a). Specifically, high inter-subject correlation (ISC) was observed in the visual cortices, spanning from the primary visual cortex to extrastriate areas and the ventral temporal regions (fusiform gyrus; Brodmann area (BA) 37, areas associated with the processing of facial information [30, 31]). Robust synchrony was also evident in the primary auditory cortices and language-associated areas (Wernicke’s area; BA 22, 39), aligning with the processing of dialogue and acoustic landscapes. In contrast, ISC was markedly lower or absent in the primary motor cortices and high-order association areas, especially the prefrontal cortex. Network-level analysis using the Yeo 17-network parcellation further confirmed this sensory-dominant pattern, with significant voxels predominantly localized within the Visual (48.67%) and Temporoparietal (11.90%) networks (Fig. 2b).

**Fig. 2.**
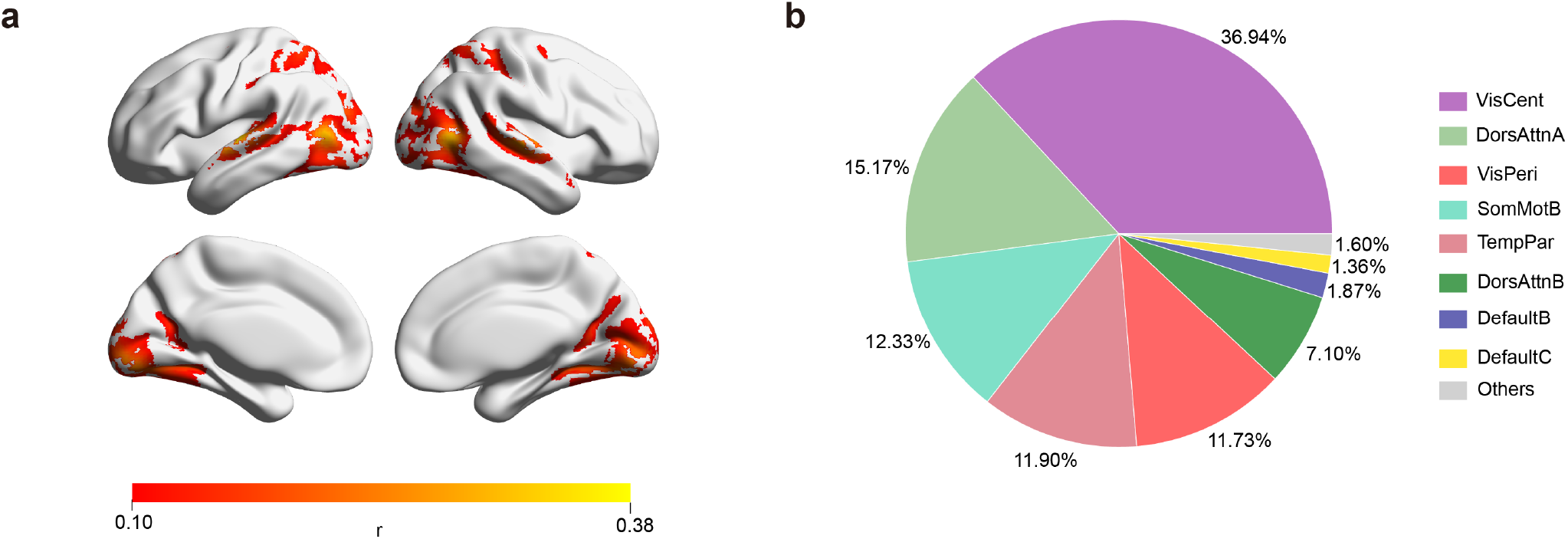
Inter-subject neural synchrony elicited by emotional movie watching. **a** Voxel-wise ISC map projected onto an inflated cortical surface, reflecting stimulus-driven synchrony across all participants (*n =* 72). High ISC is concentrated in primary visual and auditory cortices and extends into high-order perceptual regions including the fusiform gyrus and superior temporal gyrus. Color bar indicates Pearson’s *r*. **b** Network-level distribution of significant voxels across the Yeo 17-network parcellation. Networks each accounting for less than 1% of significant voxels are grouped as Others. Results are displayed at FDR-corrected *q*< 0.001with a minimum threshold of *r* > 0.10. VisCent, central visual; VisPeri, peripheral visual; DorsAttn, dorsal attention; SomMot, somatomotor; TempPar, temporoparietal; Cont, control.

### High-order prefrontal regions exhibit group-specific neural polarization

Building on the robust sensory synchrony evoked by the film, we next asked whether neural responses exhibited diagnostic-group-dependent divergence. To address this question, we quantified neural polarization by comparing within-group and between-group inter-subject synchrony (*ISC*_within–group_ − *ISC*_Between– group_ ; *Methods*) across the brain. Significant polarization was localized to two discrete voxel clusters within high-order prefrontal association cortex.

The first cluster (618 voxels; Fig. 3a) encompassed the dorsomedial prefrontal cortex (dmPFC; primarily medial BA 9) and the rostral anterior cingulate cortex (rACC; primarily pregenual BA 32). These regions are critical for emotional processing and regulation [32-34], with the dmPFC additionally specialized in tracking the interpretation of narratives [18-20, 24]. The second cluster (142 voxels; Fig. 3b) was localized in the left dorsolateral prefrontal cortex (dlPFC; primarily dorsal BA 46/9), a hub traditionally associated with executive control and cognitive regulation [35, 36]. These two clusters are hereafter referred to as the dmPFC-rACC and left dlPFC, respectively (see Supplementary Table S1 and S2 for detailed anatomical localization).

**Fig. 3.**
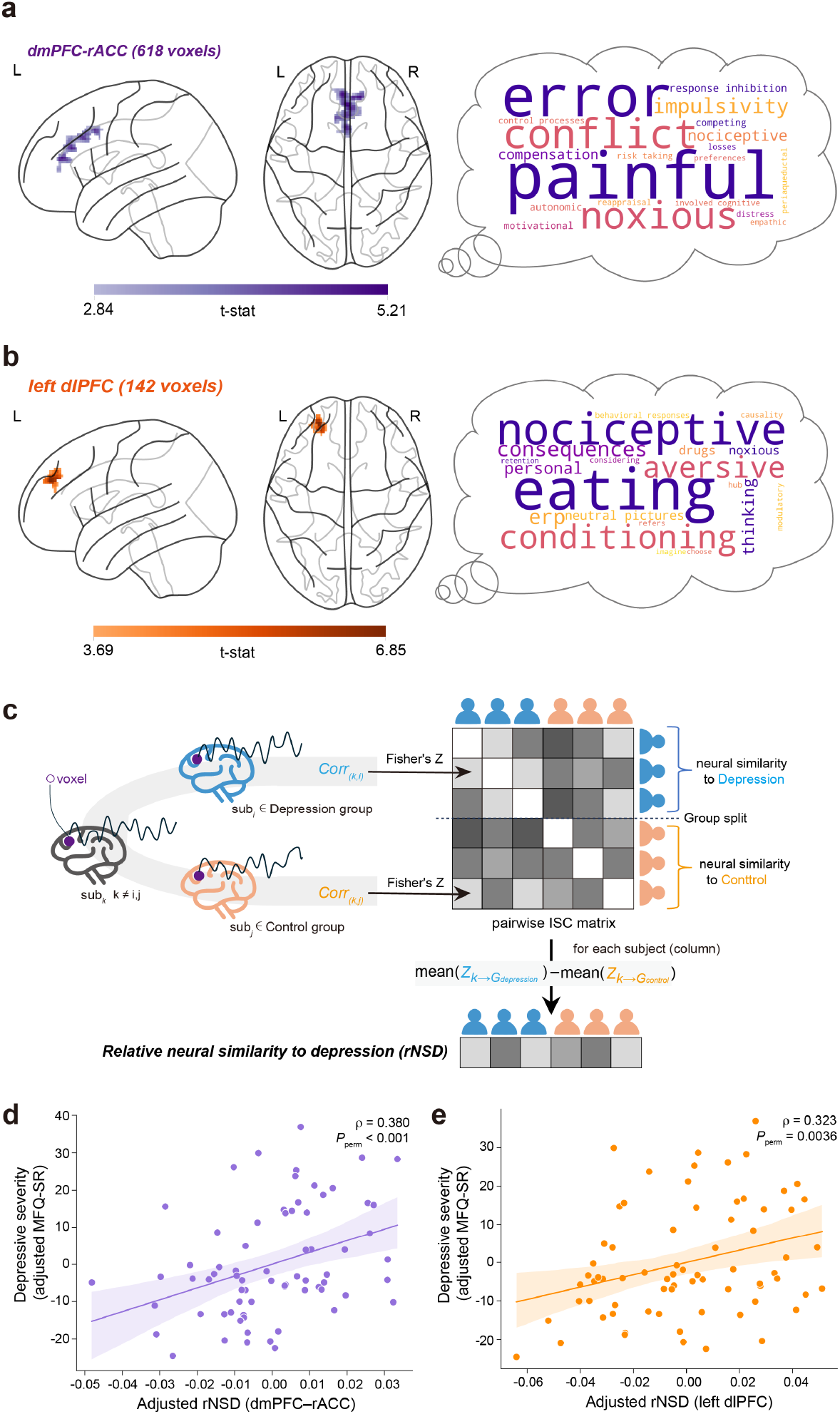
Group-specific neural polarization in high-order prefrontal regions and its association with depressive symptom severity. **a, b** Glass-brain maps (sagittal and axial views) showing voxels with significant neural polarization. Two discrete clusters were identified: (**a**) the dmPFC-rACC and (**b**) the left dlPFC. Color bars indicate the voxel-wise group-level *t*-statistic. Adjacent to each brain map, word clouds display the top 20 function-related terms derived from meta-analytic functional decoding of the corresponding spatial mask (Neurosynth χ^2^ method), with font size scaled by reverse-inference Z-score; complete rankings of the top 70 terms are provided in Supplementary Tables S3–S4. **c** Schematic illustrating the derivation of the relative neural similarity to depression (rNSD) index (see *Methods*). **d, e** Scatterplots showing the association between rNSD scores and depressive symptom severity across the entire cohort (*N=* 72). Significant positive correlations were observed for (**d**) dmPFC-rACC (partial Spearman’s ρ= 0.380, *P*_*perm*_= 0.0006) and (**e**) left dlPFC (ρ= 0.323, *P*_*perm*_ = 0.0036). Partial correlations control for age, sex, and mean framewise displacement; significance was determined by nonparametric permutation tests (10,000 iterations). Shaded bands indicate 95% confidence intervals. MFQ-SR, Moods and Feelings Questionnaire (self-report).

We further confirmed that the neural polarizations in the two clusters were not driven by a single group, as both cohorts exhibited significantly higher within-group than between-group synchrony (*p* < 0.05, Supplementary Fig. S1). To complement the categorical neural polarization analysis, we additionally examined whether similarity in depressive symptom severity was associated with similarity in neural responses across participants. A whole-brain representational similarity analysis yielded partially overlapping patterns (Supplementary Fig. S2 and Supplementary Table S5).

To evaluate the clinical relevance of these polarized patterns, we calculated a relative neural similarity to depression (rNSD) index for each participant (Fig. 3c; *Methods*). Across the entire cohort, we observed significant positive correlations between symptom severity, measured by MFQ-SR, and rNSD scores for both the dmPFC-rACC (Spearman’s ρ= 0.380, *P*_*perm*_ = 0.0006; Fig. 3d) and the left dlPFC (ρ= 0.323, *P*_*perm*_ = 0.0036; Fig. 3e). Notably, these correlations were not significant when restricted to the depression group alone (Supplementary Fig. S3), suggesting that neural polarization serves as a categorical marker of the depressed state rather than a linear gauge of symptom fluctuations within the clinical population.

### Prefrontal neural polarization extends to fronto-striatal and fronto-insular circuits

To determine whether local prefrontal divergence was embedded within broader stimulus-locked coupling patterns, we conducted inter-subject functional connectivity (ISFC) analyses using the dmPFC-rACC (Seed 1) and left dlPFC (Seed 2) polarization clusters as seeds. ISFC differs from conventional within-subject functional connectivity by correlating the time course of a seed region in one participant with target-voxel time courses in other participants during the same stimulus [16, 24, 25]. This approach isolates stimulus-locked inter-regional coupling that is shared across individuals while reducing contributions from idiosyncratic intrinsic fluctuations.

For each seed-target connection, we tested for the neural polarization effect by contrasting within-group versus between-group coupling (*ISFC* _within–group_−*ISFC*_Between– group_; see *Methods*). Significant polarization was observed for both seeds after TFCE correction (*p* < 0.05; Fig. 4), indicating that depression-related divergence was not confined to local prefrontal responses but extended to broader stimulus-locked coupling patterns. The two seeds showed polarized coupling with one another, suggesting coordinated divergence across high-order prefrontal regulatory systems. The polarized coupling also spanned distributed cortical targets, including medial prefrontal and cingulate regions, bilateral dlPFC, and the inferior frontal gyrus and anterior insula, extending into orbitofrontal cortex. Within fronto-striatal circuits, the dmPFC-rACC coupled primarily with the left ventral striatum (ventral caudate and nucleus accumbens), whereas the left dlPFC showed broader bilateral coupling across dorsal and ventral striatal divisions. The left dlPFC additionally showed selective polarized coupling with the inferior parietal lobule (IPL).

**Fig. 4.**
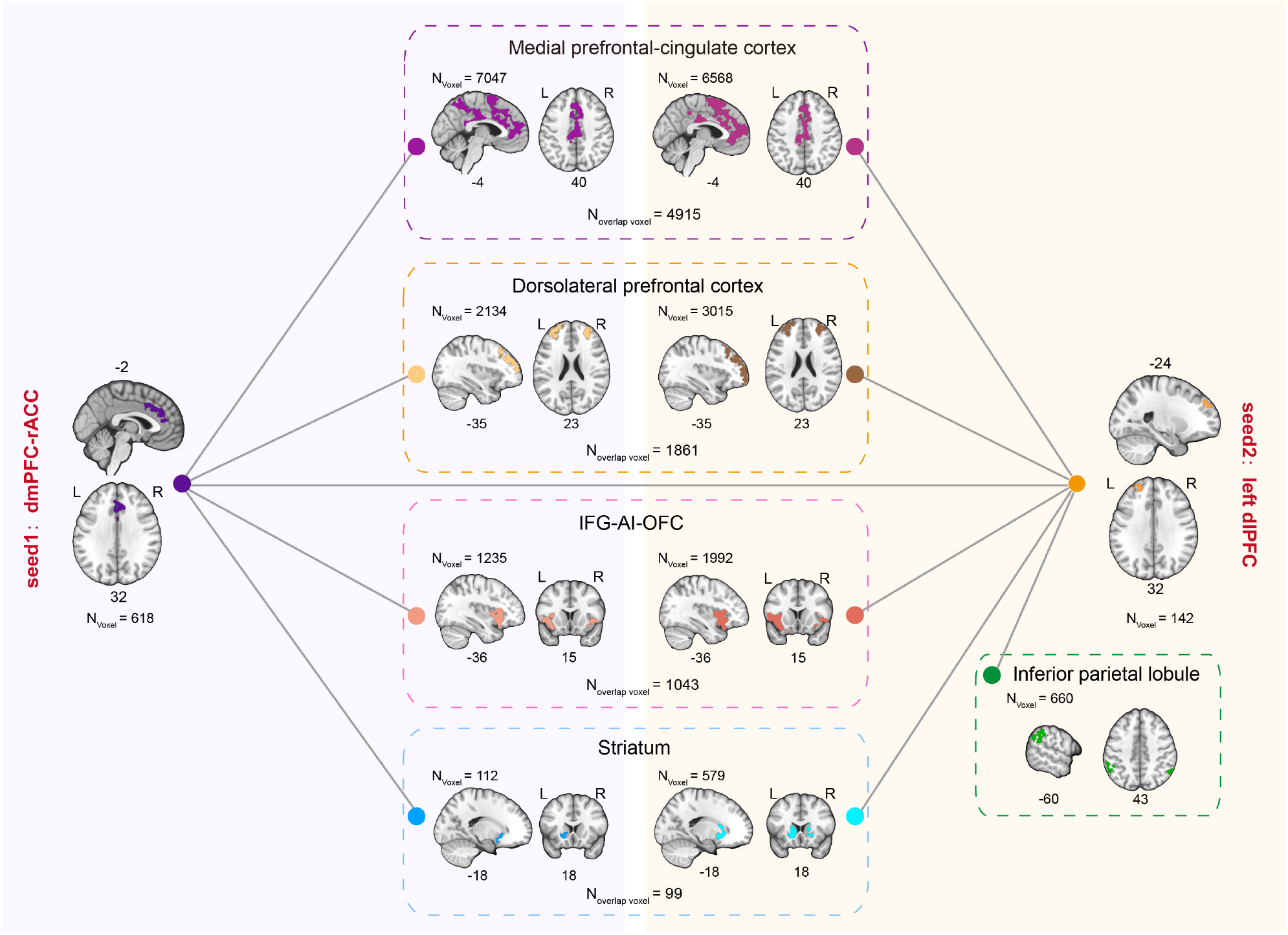
Polarized inter-subject functional connectivity linking prefrontal hubs to distributed cortical and striatal circuits. Anatomical mapping of brain regions exhibiting significantly stronger within-group than between-group stimulus-driven coupling with each seed are shown: dmPFC-rACC (Seed 1, purple) and left dlPFC (Seed 2, orange). The whole-brain ISFC divergence results are organized by approximate anatomical grouping. For each region, brain maps from Seed 1 (left) and Seed 2 (right) are displayed side by side, with voxel counts indicating the seed-specific volume (*N*_voxel_) and the spatial overlap between the two patterns (*N*_overlap_). Both seeds showed polarized coupling across a shared anatomical scaffold comprising the medial prefrontal and cingulate cortex, bilateral dlPFC, the IFG-OrG-anterior insula complex, and the striatum. The left dlPFC additionally demonstrated unique polarized coupling with the inferior parietal lobule (IPL; green dashed border). ISFC, inter-subject functional connectivity; IFG, inferior frontal gyrus; OrG, orbital gyrus.

The two seed-based maps showed substantial spatial overlap (Dice = 0.69; 8,071 common voxels), suggesting that the dmPFC-rACC and left dlPFC were embedded within a partially shared polarized coupling scaffold. Detailed voxel-level cluster statistics and Brainnetome atlas annotations are provided in Supplementary Tables S6-7. Collectively, these findings indicate that neural polarization in depression is not confined to localized prefrontal activity. Instead, it involves a systemic reorganization across the triple-network system (default mode network, central executive network, salience network) and its integrated subcortical reward circuits.

### Dynamic neural polarization selectively emerges during specific cinematic episodes

Finally, we examined the temporal dynamics of group-level divergence within the dmPFC-rACC and left dlPFC hubs to determine whether neural polarization reflected a continuously expressed property or transient divergence selectively evoked by specific narrative contexts. To this end, we quantified dynamic neural polarization magnitude (NPM), a time-resolved metric representing the summed intensity of voxels exhibiting significant polarization at each frame (*ISPS*_*within–group*_ > *ISPS*_*Between– group*_, frame-wise corrected *p* < 0.05 ; see *Methods*). The resulting NPM trajectories revealed highly dynamic and stimulus-dependent polarization patterns, with the two hubs exhibiting distinct temporal sensitivity profiles across the evolving movie narrative (Fig. 5, see Supplementary Methods for clip annotation procedure). To minimize potential instability during scan adaptation, the initial segments (Clip 1 and Clip 10) were excluded from subsequent analysis.

**Fig. 5.**
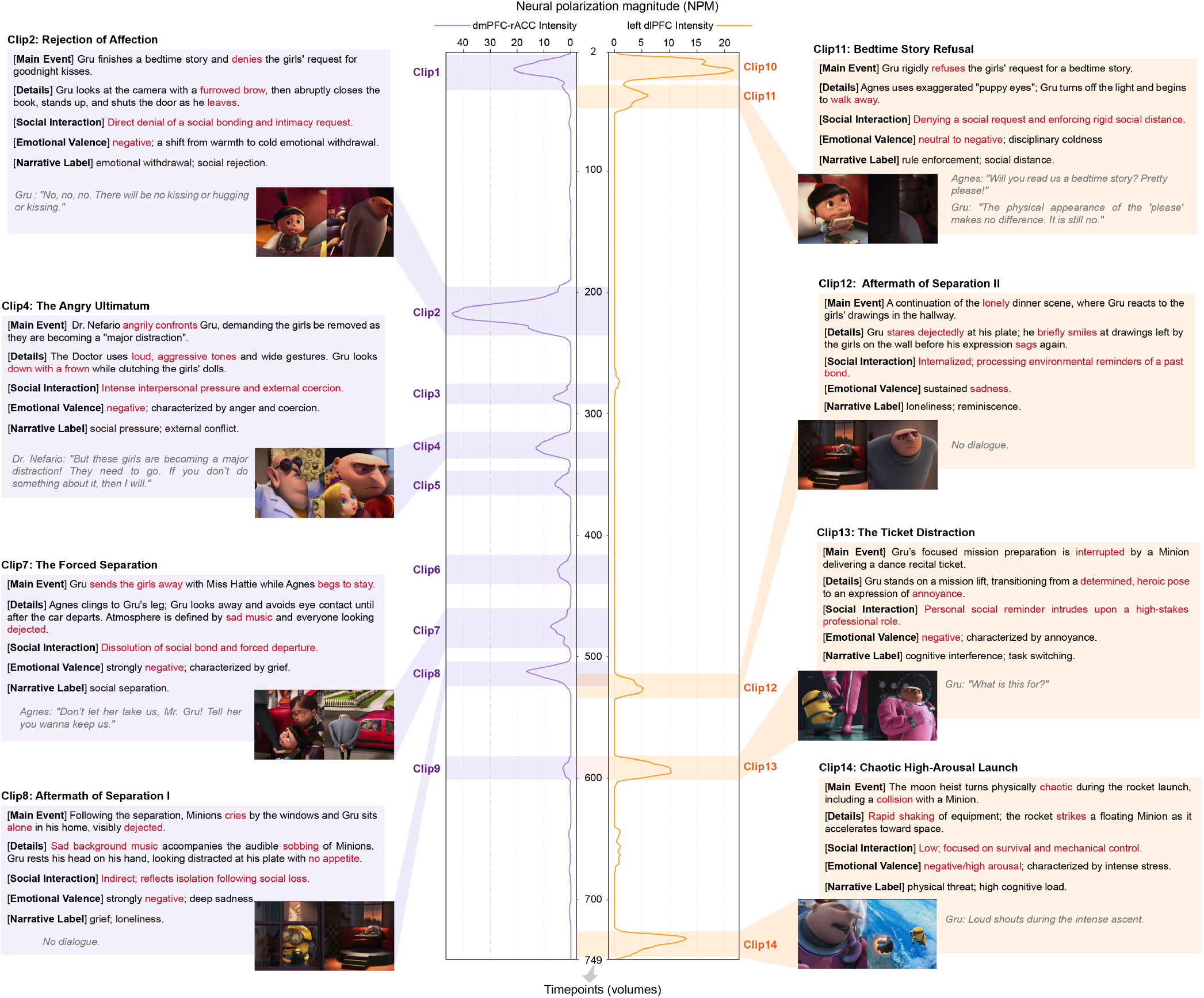
Context-specific dynamic neural polarization dissociates the dmPFC-rACC and left dlPFC. The central panel displays neural polarization magnitude (NPM) across scan time (TR 2–749) for the dmPFC-rACC (purple, left axis) and left dlPFC (orange, right axis) hubs. The trajectory is oriented with time on the vertical axis to facilitate direct alignment with the adjacent movie annotations. Shaded bands indicate significant temporal clusters. Left panels (purple): primary NPM peaks in the dmPFC-rACC, driven by socio-emotional stressors, including the Rejection of Affection (Clip2), Angry Ultimatum (Clip4), Forced Social Separation (Clip7), and Aftermath of Separation I (Clip8). Right panels (orange): NPM peaks in the left dlPFC, including lower-intensity socio-emotional episodes (Bedtime Story Refusal, Clip 11; Aftermath of Separation II, Clip 12) and high-intensity executive demand episodes (Ticket Distraction, Clip 13; Chaotic High-Arousal Launch, Clip 14). Each clip panel provides the primary cinematic event, social interaction type, emotional valence, and a representative still frame. TR, repetition time.

In the dmPFC-rACC, polarization peaks emerged predominantly during socio-affective narrative contexts involving interpersonal rejection, emotional conflict, social separation, and grief-related processing. Specifically, NPM increased sharply during scenes of the protagonist’s explicit rejection of a child’s bid for affection (Clip 2) and the aggressive ultimatum delivered by a collaborator (Clip 4). Additional polarization peaks were observed during the forced social separation of the children (Clip 7) and the subsequent non-verbal scene depicting profound solitary grief (Clip 8). Beyond these interpersonal triggers, another peak occurred during the abrupt transition from emotional tension to absurdist humor (Clip 5), suggesting that the dmPFC-rACC also tracked major affective shifts in the environmental context. Other significant episodes within this hub exhibited similar patterns of socio-emotional sensitivity (see Supplementary Fig. S4).

In contrast, the left dlPFC exhibited a divergent sensitivity profile characterized by narrative contexts related to procedural execution and cognitive control. Although polarization was observed during scenes involving request refusal (Clip 11) and solitary reminiscence (Clip 12), the NPM intensity during these socio-emotional segments remained weaker than during episodes imposing executive challenges. Specifically, polarization in the left dlPFC was strongly driven by high-stakes executive demands, peaking during sequences involving task disruption and identity conflict (Clip 13), as well as during the rocket launch scenes (Clip 14), which combined rapidly evolving threats with sustained behavioral demands.

Although most polarization peaks in the two hubs occurred at distinct time points, two specific segments revealed relatively coordinated polarization responses across the two clusters. Following the children’s departure, polarization in the dmPFC-rACC emerged immediately during the initial social loss (Clip 8), whereas the corresponding polarization peak in the left dlPFC appeared approximately 8 seconds later (Clip 12). Moreover, during the mission-interruption segment (Clip 9 and Clip 13), both hubs exhibited concurrent polarization, although the NPM was greater in the left dlPFC than in the dmPFC-rACC. Together, these findings suggest that distinct forms of narrative context are associated with different patterns of prefrontal neural divergence, with socio-emotional episodes preferentially engaging earlier dmPFC-rACC polarization and cognitively demanding episodes eliciting stronger concurrent involvement of the left dlPFC.

## Discussion

A central question in depression neuroscience is whether neural abnormalities reflect stable circuit-level dysfunctions or context-dependent states that emerge during particular forms of ongoing experience. Using naturalistic film watching, we identified a depression-related neural signature that is anatomically organized, network-embedded and dynamically gated by narrative context. Across 36 adolescents with depression and 36 matched controls, neural polarization, defined as greater within-group than between-group inter-subject synchrony, localized to two prefrontal hubs, the dmPFC–rACC and the left dlPFC. These polarized responses were clinically relevant, as individual neural similarity to the depression group scaled with depressive symptom severity across the cohort. Seed-based inter-subject functional connectivity further showed that local prefrontal divergence was embedded within broader stimulus-locked coupling patterns involving the triple-network system and its integrated subcortical reward circuits, indicating a system-level redistribution of inter-subject coordination. Most importantly, time-resolved analyses revealed that polarization was not continuously expressed throughout the film, but emerged selectively during specific narrative contexts. The dmPFC–rACC showed peak divergence during socio-emotional events such as social rejection and grief, whereas the left dlPFC showed stronger divergence during episodes of high cognitive demand. Together, these findings suggest that depression-related neural divergence during naturalistic experience is not a uniform trait-like abnormality, but a content-sensitive and state-dependent pattern of brain response.

Most neuroimaging studies of depression characterize abnormalities as task-averaged activation differences or resting-state connectivity alterations. Such approaches have been essential for identifying canonical depression-related circuits, but they provide limited insight into when these circuits diverge during ongoing experience. Our findings extend this literature by showing that prefrontal abnormalities implicated in depression are not merely present as static regional or network features. Instead, they become differentially expressed as adolescents encounter specific forms of narrative content. This distinction is particularly important for adolescence, when social evaluation, interpersonal loss, affective instability, and developing cognitive control systems are central to depression vulnerability [37-39]. Naturalistic stimulation therefore provides a setting in which depression-related circuit abnormalities can be examined not only as stable traits, but also as temporally specific neural states elicited by meaningful social and emotional contexts.

The spatial distribution of polarization links depression-related divergence to prefrontal systems that are both clinically relevant and well suited to naturalistic social-emotional processing. The medial cluster, dmPFC-rACC, dominated by rostral cingulate subdivisions and dmPFC extension, has been repeatedly implicated in depression, including abnormalities in default-mode function, rumination, self-referential processing, and affective regulation [9, 32, 34, 40-43]. Moreover, dmPFC also tracks narrative interpretation and inferences about other minds [18-20]. The dmPFC-rACC polarization may therefore reflect diagnostic-group differences in how adolescents interpret and regulate socially meaningful narrative content. The left dlPFC cluster, by contrast, falls within a lateral prefrontal control system implicated in cognitive control and the selection of emotion-regulation strategies, both of which are disrupted in depression [35, 36, 44, 45]. Thus, the two polarized hubs do not merely reproduce known depression-related regions. They identify medial and lateral prefrontal systems through which diagnostic-group divergence becomes expressed during naturalistic emotional experience.

Notably, polarization at the left dlPFC was driven asymmetrically by the depressed group. Although within-group synchrony exceeded between-group synchrony in both groups, the magnitude of this within–between difference was substantially larger in depressed adolescents than in controls (Supplementary Results and Supplementary Fig. S1). This asymmetry is consistent with cognitive inflexibility, a core depression vulnerability characterized by reduced capacity to engage varied cognitive strategies under changing affective demands [46]. The pattern aligns with behavioral evidence of impaired cognitive control over negative material [41, 42]. Neural homogenization at this cognitive-control hub may parallel depression’s behavioral inflexibility, extending it into how the brain engages with affective content.

Polarization at these two hubs is not a localized phenomenon, but embedded within broader stimulus-locked coupling patterns. Seed-based ISFC revealed that both the dmPFC-rACC and left dlPFC showed polarized coupling with distributed cortical and subcortical targets, including additional medial prefrontal and cingulate regions, bilateral dorsolateral prefrontal regions, inferior parietal cortex, insular cortex and striatal circuitry. The two seeds anchored a substantially overlapping scaffold (Dice = 0.69), pointing to a shared network-level signature rather than two hub-specific dysfunctions. The hubs were also polarized with one another, a stimulus-evoked instance of the executive–affective coordination failure that cognitive theories of depression have long emphasized [42]. This network-embedded pattern complements views of depression as a disorder of large-scale brain organization, particularly involving default-mode, salience, frontoparietal control, and fronto-striatal systems [8, 47, 48]. However, the present findings differ from conventional resting-state connectivity results in an important respect. ISFC captures stimulus-locked inter-regional coupling across participants exposed to the same stimulus, rather than spontaneous within-subject connectivity. Thus, our results suggest that depression-related divergence is expressed not only in local prefrontal responses, but also in how these prefrontal systems participate in shared stimulus-locked coupling with affective, control, salience, and reward-related circuits during ongoing experience.

Crucially, the temporal profiles of these hubs further clarified their functional relevance, revealing that polarization was not a constant feature of the depressed brain but emerged transiently, in response to specific events in the film. The dmPFC–rACC carried polarization most strongly during scenes of negative interpersonal exchange and emotional distress: the protagonist’s rejection of a child’s bid for affection, an aggressive ultimatum from a collaborator, the forced separation of the children, and the prolonged depiction of solitary grief that followed. An additional peak coincided with an abrupt narrative shift from emotional tension to absurdist humor, suggesting that the hub also tracked major affective transitions. At the dmPFC–rACC, divergence concentrated in the affective situations long known to matter most for the disorder, namely interpersonal injury, loss, and abrupt affective change [49, 50]. These contexts align closely with interpersonal and affective domains that are central to adolescent depression, but the present data add temporal specificity by showing when such divergence becomes most pronounced during an unfolding stimulus.

The left dlPFC’s content profile diverged sharply. Its polarization peaks were anchored in episodes of high cognitive demand: the task-switching triggered by an unexpected interruption to a high-stakes mission, and a chaotic rocket-launch sequence dominated by physical threat and rapid information processing. During these mission-interference scenes, both hubs polarized concurrently, but the left dlPFC dominated, consistent with the increased executive load. A more noteworthy pattern emerged in the aftermath of the children’s departure. The dmPFC–rACC polarized immediately at the onset of social loss, whereas the left dlPFC’s polarization followed approximately eight seconds later. This sequence is consistent with the broader principle that initial affective processing precedes downstream cognitive engagement in emotion regulation [35]. Across these temporal profiles, the dmPFC–rACC and left dlPFC each carried depression’s stimulus-evoked divergence at different moments and in response to different facets of the unfolding stimulus. Together, these findings support the view that depression-related neural abnormalities are not uniformly expressed, but selectively emerge when ongoing experience engages particular prefrontal systems.

The present findings also raise cautious clinical and translational implications. First, the rNSD results suggest that polarization may primarily reflect diagnostic-group organization of neural responses rather than continuous symptom variation within the depressed group. Across the full cohort, individuals whose responses at the two hubs were more similar to the typical depression-group pattern reported greater depressive symptoms, but this association did not hold within the depression group alone. Second, the content specificity of polarization may help characterize the experiential contexts in which depression-related neural divergence becomes most apparent. The strongest polarization episodes were linked to recognizable narrative contexts, including interpersonal rejection, separation, grief, abrupt affective transitions, procedural disruption, and rapidly changing task demands. Such context-sensitive signatures may provide a bridge between naturalistic neuroimaging and clinically meaningful situations encountered in daily life, although their relevance for psychoeducation or symptom management requires direct behavioral and clinical validation. Finally, these findings suggest a speculative but testable implication for state-dependent intervention. Both polarized hubs overlap systems already relevant to clinical neuromodulation in depression, including cingulate targets for deep brain stimulation [51] and left dlPFC targets for repetitive transcranial magnetic stimulation [52]. Current neuromodulation strategies primarily emphasize anatomical target selection, but our results raise the possibility that the neural state induced by ongoing emotional or cognitive context may also influence treatment response. Each hub’s polarization concentrates in moments of content matched to its functional profile. Socio-emotional events drove the dmPFC–rACC, whereas episodes of cognitive demand drove the left dlPFC. This raises the possibility that stimulation could be more effective when delivered while the relevant circuitry is actively engaged with such content. This conjecture is plainly speculative. But the dynamic, content-specific structure of polarization revealed here suggests that depression’s divergence is not equally accessible at all moments, and that the timing of intervention may merit more direct investigation.

A methodological implication of this work is that signal-driven temporal localization can complement annotation-driven approaches in naturalistic neuroimaging [11]. Prior studies often begin with experimenter-defined stimulus features, such as continuous valence or emotional intensity ratings, semantic labels, or positive and negative movie segments, and then test how neural activity tracks those features [22, 24, 25]. Our framework reverses this logic. Voxel-wise polarization based on inter-subject phase synchrony [27] first identifies the moments at which neural divergence between groups is statistically significant, and the cinematic content of those moments is then examined post hoc. This offers two methodological advantages. First, the temporal grain of detection is set by the signal rather than by the stimulus annotation, allowing per-TR localization that sliding-window approaches cannot resolve. Second, the framework requires no a priori commitments about which stimulus features matter, nor any assumption that ratings from independent annotators reflect the affective experience of the patient population. It identifies behaviorally meaningful content from any naturalistic stimulus the brain reacts to differentially, expanding the range of materials that can be productively analyzed for group-specific neural patterns. The approach is hypothesis-generating in spirit, complementing the hypothesis-testing logic of feature-driven analyses, and can be deployed across disorders and stimulus types to identify the specific moments at which group-level neural divergence emerges. In this sense, signal-driven polarization can generate candidate moments of clinical relevance, while quantitative content annotations can test which stimulus features account for their timing.

Several limitations bound the inferences we can draw, and each points to a productive next step. First, although polarization moments were identified directly from neural signal, the cinematic content of those moments was annotated qualitatively. Pairing signal-driven temporal localization with quantitative content features drawn from vision and language models could provide continuous, scalable maps of content-brain relationships. Second, our study deliberately focused on adolescents, a developmental period in which depression often emerges and socio-emotional processing is particularly salient. The findings should therefore be interpreted primarily in the context of adolescent depression. Future work in younger children, adults, and longitudinal developmental cohorts will be needed to determine whether the observed polarization patterns are specific to adolescence or reflect more general features of depression across development. Third, the temporal resolution of BOLD fMRI and hemodynamic variability limit strong claims about precise processing sequences, including short latency differences between hubs. Finally, complementary cluster-level analyses recovered only a subset of the narrative-associated polarization episodes identified by voxel-wise NPM analyses (Supplementary Results, Supplementary Figs. S5-6). This likely reflects reduced sensitivity to spatially focal and transient effects after cluster-level averaging, and suggests that dynamic polarization during naturalistic experience may be spatially constrained rather than uniformly expressed across entire clusters.

Taken together, this work reframes depression-related neural signatures as dynamic and context-dependent patterns of brain response during ongoing experience. By identifying where group divergence localizes, how it is embedded within stimulus-locked coupling networks, and when it emerges during specific narrative contexts, the framework moves beyond static descriptions of depression-related circuitry. More broadly, it suggests that psychiatric neural differences may become most informative when they are studied not only as properties of individuals or circuits, but also as states elicited by the changing structure of real-world experience.

## Methods

### Participants

Data for this study were obtained from the Child Mind Institute’s Healthy Brain Network (HBN) portal [53], a large-scale, ongoing initiative recruiting a high-risk community sample of children and adolescents with various clinical concerns. Written informed consent was obtained from all participants; for those under 18 years of age, a parent or legal guardian provided written consent and the participant provided written assent. The HBN project was approved by the Chesapeake Institutional Review Board.

Imaging and phenotypic data were downloaded for 221 unmedicated participants (aged 16–21 years) with complete demographic records (age, sex, and scanning site) from HBN Releases 1–9. Of these, 143 had complete functional MRI (fMRI) data for the *Despicable Me* movie-watching paradigm and passed the standardized preprocessing pipeline. Five participants were excluded for excessive head motion (mean framewise displacement, FD ≥ 0.5 mm), yielding 138 eligible participants, of whom 36 carried a clinical diagnosis of depression. Clinical diagnoses are assigned by the clinician after administering the Kiddie Schedule for Affective Disorders and Schizophrenia (KSADS) [54] and considering additional data and interactions provided in the participation.

To ensure a well-balanced group comparison, a propensity-score matching analysis was performed using the MatchIt package in R. Matching criteria included age, sex, and five psychiatric comorbidities: Generalized Anxiety Disorder, Social Anxiety Disorder, Autism Spectrum Disorder, and Attention-Deficit/Hyperactivity Disorder (Inattentive and Combined subtypes). This procedure yielded a final analytical sample of 72 participants: a depression group (n = 36) and a matched control group (n = 36). Specifically, the depression group included 20 males and 16 females (mean age = 17.70 ± 1.43 years), with the control group consisting of 17 males and 19 females (mean age = 18.16 ± 1.71 years). Self-report responses to the Moods and Feelings Questionnaire (MFQ-SR) were used to quantify depressive symptom severity. The MFQ-SR comprises 33 statements describing affective states over the preceding two weeks (e.g., ‘I felt miserable or unhappy’; ‘I felt lonely’), each rated on a 3-point scale (not true, sometimes, true), and has been validated for use in clinical and sub-clinical pediatric populations [29]. Group differences in demographic and clinical variables were assessed using independent-samples t-tests for continuous variables and chi-square tests for categorical variables. The phenotypic variables used for participant selection are summarized in Supplementary Table S8.

### MRI Data preprocessing

Structural and functional MRI data were preprocessed using fMRIPrep [55]. The T1-weighted image was skull-stripped, and brain tissue was segmented into white matter, gray matter, and cerebrospinal fluid using FreeSurfer [56]. The T1w reference was spatially normalized to the MNI152NLin6Asym template using ANTs nonlinear registration [57]. For the fMRI, a reference BOLD volume was generated and head motion was corrected using FSL mcflirt [58]. Susceptibility distortion correction was applied using the available fieldmap. The BOLD reference was coregistered to the T1w image using boundary-based registration (BBR; 6 degrees of freedom) [59]. Confound time series, including head motion parameters (framewise displacement), anatomical and temporal CompCor components, and global signal, were estimated for subsequent denoising. The preprocessed BOLD data were resampled into MNI152NLin6Asym standard space at 2 mm isotropic resolution.

Further denoising was performed using XCP-D [60]. A 27-parameter nuisance regression strategy was applied [61, 62], comprising six rigid-body motion parameters, their temporal derivatives, and the quadratic expansion of all 12 terms, together with the mean global signal, mean white matter signal, and mean cerebrospinal fluid signal. Data were band-pass filtered using a second-order Butterworth filter (0.01–0.1 Hz). This frequency range was selected because dynamic emotional responses to naturalistic stimuli are understood to fluctuate primarily below 0.1 Hz [22, 63]. Furthermore, data were spatially smoothed with a Gaussian kernel (FWHM = 6 mm). Prior to all subsequent analyses, each voxel’s time series was z-scored across time to standardize signal amplitude across participants and scanning sites.

### Static inter-subject correlation analyses

Inter-subject correlation (ISC) analysis was performed using a pairwise approach to quantify stimulus-driven neural synchrony across participants [14, 20, 25, 63, 64]. For each voxel, the Pearson correlation coefficient between the BOLD time courses of every participant pair (*i, j*) was computed and Fisher *z*-transformed to ensure statistical normality [65-67], yielding a symmetric subject-by-subject correlation matrix per voxel.

#### Shared Neural Response Analysis

To characterize the stimulus-evoked synchrony shared across the full cohort, a subject-specific ISC value was computed for each voxel (Fig. 1b(1)). For a given participant *i*, this value was the average of the Fisher *z* -transformed pairwise correlations between participant *i* and all remaining participants (*j* ≠ *i*). These per-participant maps were averaged across all participants to produce a group-level mean Fisher *z*-map, subsequently back-transformed to an *r*-map for visualization. Statistical significance was assessed using a nonparametric permutation test [24, 25, 67]. At each voxel, the observed *t*-statistic tested whether the mean Fisher *z*-value exceeded zero. A null distribution was constructed by randomly sign-flipping the Fisher *z*-values of a random subset of participants and recomputing the one-sample *t*-statistic (10,000 permutations). The *P*-value was the proportion of null statistics exceeding the observed value. Voxel-wise multiple comparisons were corrected using the Benjamini–Hochberg false discovery rate (FDR) procedure [68] (*q*< 0.001).

#### Static Neural Polarization Analysis

To identify voxels exhibiting group-biased neural processing—where activity was more synchronized within groups than between them—we quantified a neural polarization index [24] (Fig. 1b(2)). For each participant *i* and voxel *k*, the pairwise Fisher *z*-values were partitioned into the within-group ISC 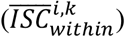, defined as the mean correlation between participant *i* and all other members of the same diagnostic group, and the between-group ISC 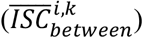, defined as the mean correlation between participant *i* and all members of the other group. The neural polarization index was computed as:

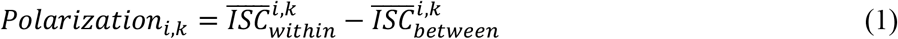

A positive value indicates that a participant’s neural response pattern is more similar to their own group than to the other. Subject-level polarization maps were averaged across all participants to obtain a group-level mean difference map (in Fisher *z*-values). Statistical significance was assessed via a voxel-wise one-sample *t*-test against zero (one-tailed; testing *ISC*_*within*_ > *ISC*_*between*_), with multiple comparisons corrected using threshold-free cluster enhancement (TFCE) [69]. TFCE was selected for its favorable balance between family-wise error rate control and test-retest reliability [70]. Permutation testing (10,000 iterations) was implemented in the DPABI toolbox [71, 72], with a significance threshold of *P* < 0.05.

#### Relative Neural Similarity to Depression (rNSD) Analysis

To quantify how closely each participant’s neural response resembled the depression group relative to the control group, we computed an rNSD index (Fig. 3c). For each participant *i* and voxel *k*, we derived the similarity to the depression group 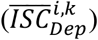, defined as the mean correlation between participant *i* and all members of the depression group, and the similarity to the control group 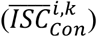, defined as the mean correlation between participant *i* and all members of the control group. The rNSD value was computed as:

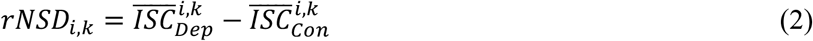

This analysis was restricted to voxels within the two ROIs that were identified by the static neural polarization analysis (dmPFC-rACC and left dlPFC). A single subject-level rNSD score per ROI was obtained by averaging voxel-wise values across all voxels in the region. The association between rNSD scores and self-reported depressive symptom severity (MFQ-SR) was assessed using partial Spearman rank correlations, controlling for age, sex, and mean framewise displacement [22]. Statistical significance was evaluated via a nonparametric permutation test, which preserved the distributional characteristics of the data while destroying the specific link between the neural and behavioral variables. A null distribution of the partial Spearman correlation was generated by randomly shuffling the rNSD scores across participants (10,000 permutations) while keeping MFQ-SR scores and covariates fixed. The *P*-value was the proportion of permuted correlations equal to or more extreme than the observed value (*P* < 0.05, one-tailed).

### Inter-subject functional connectivity analyses

To examine whether the identified local neural polarization in high-order prefrontal regions reflects a broader reorganization of stimulus-driven network communication, we performed inter-subject functional connectivity (ISFC) analyses [16, 73]. Unlike standard functional connectivity, ISFC measures stimulus-evoked communication across brain regions and can reveal stimulus-related functional networks [63]. Specifically, for each participant pair (*i, j*), ISFC was computed by correlating the time course of a seed region in participant *i* with the target time course in participant *j*. The two regions exhibiting significant static neural polarization, dmPFC-rACC (Seed 1) and left dlPFC (Seed 2), served as seed regions. For each participant, the BOLD time course was extracted and averaged across all voxels within each seed region.

*ISFC divergence*. Group differences in inter-subject functional coupling were assessed using the ISFC divergence metric [24] (Fig. 1b(3)). For each connection (seed-to-seed or seed-to-voxel), the pairwise ISFC estimates described above were Fisher *z* -transformed and partitioned by group membership. The ISFC divergence for participant *i* was defined as:

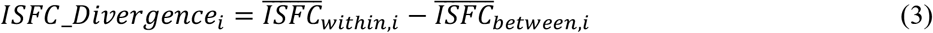

where 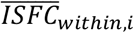 and 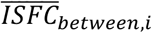 denote the mean Fisher *z*-transformed coupling between participant *i* and all within-group and between-group members, respectively.

*Seed-to-seed functional connectivity*. We examined the inter-subject functional coupling between the two seed regions firstly. For each participant, the ISFC divergence was calculated between the time courses of dmPFC-rACC and left dlPFC. Statistical significance was assessed using a nonparametric permutation test: a null distribution was generated by randomly sign-flipping the divergence values across participants and recomputing the group-level *t*-statistic (10,000 permutations). The *P*-value was the proportion of null statistics exceeding the observed value (*P* < 0.05, one-tailed).

#### Seed-to-whole-brain functional connectivity

To characterize the full network extent of polarized functional coupling, we performed a seed-to-voxel ISFC analysis for each seed region. The ISFC divergence was computed between the seed time course and the time course of every other voxel in the brain (excluding voxels within either seed region), yielding a whole-brain divergence map per participant. Statistical inference followed the same procedure as the static neural polarization analysis: voxel-wise one-sample *t*-tests (one-tailed) corrected for multiple comparisons using TFCE [69] with 10,000 permutation iterations as implemented in DPABI [71, 72], at a significance threshold of *P* < 0.05.

### Dynamic neural polarization analysis

To characterize the temporal dynamics of group-level neural divergence, we employed inter-subject phase synchrony (ISPS) analysis [27] (Fig. 1b(4)). Unlike Pearson correlation, which yields a single estimate over the entire time course, ISPS quantifies instantaneous synchrony by extracting the phase of the signal independently of its amplitude.

For each voxel within the two ROIs (dmPFC-rACC and left dlPFC), the instantaneous phase *ϕ*(*t*) of the BOLD time series was extracted via the Hilbert transform, converting the real-valued signal into an analytic signal. To minimize edge artifacts introduced by the transform, the first and last time points were discarded, yielding 748 effective time frames for subsequent analysis. Pairwise phase synchrony between participants *i* and *j* at each time point *t* was computed as the cosine of their instantaneous phase difference:

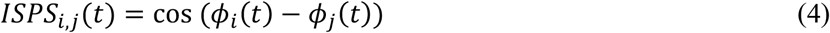

This metric ranges from −1 (anti-phase synchronization) to 1 (perfect in-phase synchronization). All values were Fisher *z*-transformed to normalize the distribution for statistical testing.

#### Voxel-wise estimation of dynamic neural polarization

For each participant *i* and each time point *t*, the dynamic neural polarization was computed as the difference between the average within-group and between-group ISPS:

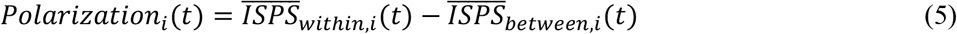

This yielded a time-resolved polarization trajectory for each voxel of each participant. Significant temporal windows of group divergence were identified using a nonparametric temporal cluster-based permutation test [74, 75], as implemented in the MNE-Python package. At each time point, a one-sample *t*-test assessed whether the polarization index exceeded zero. Consecutive time points surpassing an uncorrected threshold of *P* < 0.05 were grouped into temporal clusters, and each cluster’s statistic was defined as the sum of its constituent *t*-values. A null distribution of maximal cluster statistics was generated by randomly sign-flipping the polarization time courses across participants (10,000 permutations). Temporal clusters were considered significant if their mass exceeded the 95th percentile of the null distribution (*P* < 0.05, one-tailed, corrected).

#### Neural polarization magnitude

To summarize regional dynamics as a single time-resolved index, we computed a neural polarization magnitude (NPM) trajectory for each ROI. At each time point *t*, the NPM was defined as the sum of the group-average polarization values across all voxels within the ROI that reached temporal significance:

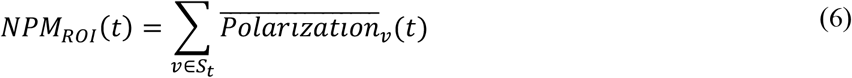

where *S*_*t*_ denotes the set of voxels within the ROI exhibiting significant polarization at time *t*. This metric integrates information about both the spatial extent and the magnitude of group divergence, providing an instantaneous index of regional neural polarization.

To align NPM peaks with specific movie content, hemodynamic response delay was accounted for by shifting the neural signal 6 TRs earlier [22]. The movie timestamp corresponding to each TR was therefore calculated as:

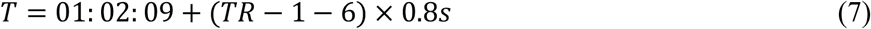

where TR = 1 corresponds to scan onset, such that TR=7 aligned with the movie clip onset (01:02:09) and the first 6 TRs were excluded from content-level interpretation.

## Code Availability

Preprocessing was performed using fMRIPrep (v 24.1.1; https://fmriprep.org) [55] and XCP-D (v 0.10.7; https://xcp-d.readthedocs.io)[60]. Voxel-wise permutation testing and TFCE-based inference for the static polarization and ISFC analyses were implemented in the DPABI toolbox (http://rfmri.org/dpabi) [71, 72]. Temporal cluster-based permutation testing for the dynamic polarization analysis used MNE-Python (‘permutation_cluster_1samp_test’; https://mne.tools) [76]. Custom analysis code for ISC, ISFC, and ISPS analyses, along with all statistical procedures described in this manuscript, is available at https://github.com/YidaoWen/movie-fMRI-dynamic-NDV-depression.

## Acknowledgments

This work was supported by the STI2030-Major Projects (Grant No. 2021ZD0200200 to T.J.), National Natural Science Foundation of China (Grant No. 62327805 to T.J., No. 62403465 to S.W.) and China Postdoctoral Science Foundation (GZC20232999 to S.W.). The analyses in this paper were made possible by the data provided by the Child Mind Institute as part of the Healthy Brain Network (HBN) Biobank. We sincerely appreciate the participants for their contributions and the HBN team for their dedication to data collection, processing, and dissemination. We would also like to thank our colleagues at the Brainnetome Center for their ongoing support and insightful discussions.

## Author Contributions

Q.L. and W.S. conceptualized the study. Q.L., H. W., and W. L. developed the methodology. Q.L., W.S. and N. L. curated the data. Q.L., W. S., H. W., W. L., N. L., S. D, C. C., and L. F. conducted the investigation. Q.L. and W.S. contributed to the visualization. T.J. supervised the study. Q.L. and W. S. drafted the original manuscript. C. C., L. F., and T.J. reviewed and edited the manuscript.

## Competing Interests

The authors declare no competing interests.

Data are presented as mean (standard deviation) for continuous variables and as count (percentage) for categorical variables. Independent-sample *t* -tests were used for age and MFQ-SR comparisons, and χ^2^ (Chi-square) tests were used for categorical variables. All tests were two-tailed. *p*-values represent the significance of the difference between the depression and control groups (depression minus Control). RUBIC, Rutgers University Brain Imaging Center; CBIC, CitiGroup Cornell Brain Imaging Center; CUNY, City College of New York; GAD, Generalized Anxiety Disorder; SAD, Social Anxiety Disorder; ASD, Autism Spectrum Disorder; ADHD, Attention-Deficit/Hyperactivity Disorder; MFQ-SR, Mood and Feelings Questionnaire-Self Report.

## Notes

### Competing Interest Statement

The authors have declared no competing interest.

## Reference

1. Gotlib, I.H. and J. Joormann, Cognition and depression: current status and future directions. Annual review of clinical psychology, 2010. 6: p. 285–312.

2. Collaborators, G.M.D., Global, regional, and national burden of 12 mental disorders in 204 countries and territories, 1990–2019: a systematic analysis for the Global Burden of Disease Study 2019. The Lancet Psychiatry, 2022. 9(2): p. 137–150.

3. Avenevoli, S., et al., Major depression in the national comorbidity survey–adolescent supplement: Prevalence, correlates, and treatment. Journal of the American Academy of Child & Adolescent Psychiatry, 2015. 54(1): p. 37–44. e2.

4. Hawton, K., K.E. Saunders, and R.C. O’Connor, Self-harm and suicide in adolescents. The lancet, 2012. 379(9834): p. 2373–2382.

5. Disner, S.G., et al., Neural mechanisms of the cognitive model of depression. Nature reviews neuroscience, 2011. 12(8): p. 467–477.

6. Siegle, G.J., et al., Can’t shake that feeling: event-related fMRI assessment of sustained amygdala activity in response to emotional information in depressed individuals. Biological psychiatry, 2002. 51(9): p. 693–707.

7. Diener, C., et al., A meta-analysis of neurofunctional imaging studies of emotion and cognition in major depression. Neuroimage, 2012. 61(3): p. 677–685.

8. Kaiser, R.H., et al., Large-scale network dysfunction in major depressive disorder: a meta-analysis of resting-state functional connectivity. JAMA psychiatry, 2015. 72(6): p. 603–611.

9. Sheline, Y.I., et al., Resting-state functional MRI in depression unmasks increased connectivity between networks via the dorsal nexus. Proceedings of the National Academy of Sciences, 2010. 107(24): p. 11020–11025.

10. Javaheripour, N., et al., Altered resting-state functional connectome in major depressive disorder: a mega-analysis from the PsyMRI consortium. Translational psychiatry, 2021. 11(1): p. 511.

11. Sonkusare, S., M. Breakspear, and C. Guo, Naturalistic stimuli in neuroscience: critically acclaimed. Trends in cognitive sciences, 2019. 23(8): p. 699–714.

12. Jääskeläinen, I.P., et al., Movies and narratives as naturalistic stimuli in neuroimaging. NeuroImage, 2021. 224: p. 117445.

13. Nastase, S.A., A. Goldstein, and U. Hasson, Keep it real: rethinking the primacy of experimental control in cognitive neuroscience. NeuroImage, 2020. 222: p. 117254.

14. Hasson, U., et al., Intersubject synchronization of cortical activity during natural vision. science, 2004. 303(5664): p. 1634–1640.

15. Hasson, U., R. Malach, and D.J. Heeger, Reliability of cortical activity during natural stimulation. Trends in cognitive sciences, 2010. 14(1): p. 40–48.

16. Simony, E., et al., Dynamic reconfiguration of the default mode network during narrative comprehension. Nature communications, 2016. 7(1): p. 12141.

17. Parkinson, C., A.M. Kleinbaum, and T. Wheatley, Similar neural responses predict friendship. Nature communications, 2018. 9(1): p. 332.

18. Yeshurun, Y., et al., Same story, different story: the neural representation of interpretive frameworks. Psychological science, 2017. 28(3): p. 307–319.

19. Nguyen, M., T. Vanderwal, and U. Hasson, Shared understanding of narratives is correlated with shared neural responses. NeuroImage, 2019. 184: p. 161–170.

20. Finn, E.S., et al., Trait paranoia shapes inter-subject synchrony in brain activity during an ambiguous social narrative. Nature Communications, 2018. 9.

21. Guo, C.C., et al., Out-of-sync: disrupted neural activity in emotional circuitry during film viewing in melancholic depression. Scientific reports, 2015. 5(1): p. 11605.

22. Gruskin, D.C., M.D. Rosenberg, and A.J. Holmes, Relationships between depressive symptoms and brain responses during emotional movie viewing emerge in adolescence. NeuroImage, 2020. 216: p. 116217.

23. Cahart, M.-S., V. Giampietro, and O. O’Daly, Atypical attentional network dynamics in adolescent depression during emotional movie viewing. Social Cognitive and Affective Neuroscience, 2025. 20(1): p. nsaf011.

24. Leong, Y.C., et al., Conservative and liberal attitudes drive polarized neural responses to political content. Proceedings of the National Academy of Sciences, 2020. 117(44): p. 27731–27739.

25. Zhang, M., et al., Neural divergence between individuals with and without minor depression during dynamic emotion processing: a movie-fMRI Study. Social Cognitive and Affective Neuroscience, 2024. 19(1): p. nsae086.

26. Hutchison, R.M., et al., Dynamic functional connectivity: promise, issues, and interpretations. Neuroimage, 2013. 80: p. 360–378.

27. Glerean, E., et al., Functional magnetic resonance imaging phase synchronization as a measure of dynamic functional connectivity. Brain connectivity, 2012. 2(2): p. 91–101.

28. Vanderwal, T., J. Eilbott, and F.X. Castellanos, Movies in the magnet: Naturalistic paradigms in developmental functional neuroimaging. Developmental cognitive neuroscience, 2019. 36: p. 100600.

29. Angold, A., et al., Development of a short questionnaire for use in epidemiological studies of depression in children and adolescents. International journal of methods in psychiatric research, 1995.

30. Kanwisher, N., J. McDermott, and M.M. Chun, The fusiform face area: a module in human extrastriate cortex specialized for face perception. Journal of neuroscience, 1997. 17(11): p. 4302–4311.

31. Haxby, J.V., E.A. Hoffman, and M.I. Gobbini, The distributed human neural system for face perception. Trends in cognitive sciences, 2000. 4(6): p. 223–233.

32. Etkin, A., T. Egner, and R. Kalisch, Emotional processing in anterior cingulate and medial prefrontal cortex. Trends in cognitive sciences, 2011. 15(2): p. 85–93.

33. Bush, G., P. Luu, and M.I. Posner, Cognitive and emotional influences in anterior cingulate cortex. Trends in cognitive sciences, 2000. 4(6): p. 215–222.

34. Phillips, M.L., et al., Neurobiology of emotion perception I: The neural basis of normal emotion perception. Biological psychiatry, 2003. 54(5): p. 504–514.

35. Ochsner, K.N. and J.J. Gross, The cognitive control of emotion. Trends in cognitive sciences, 2005. 9(5): p. 242–249.

36. Miller, E.K. and J.D. Cohen, An integrative theory of prefrontal cortex function. Annual review of neuroscience, 2001. 24(1): p. 167–202.

37. Davey, C.G., M. Yücel, and N.B. Allen, The emergence of depression in adolescence: development of the prefrontal cortex and the representation of reward. Neurosci Biobehav Rev, 2008. 32(1): p. 1–19.

38. Crone, E.A. and R.E. Dahl, Understanding adolescence as a period of social–affective engagement and goal flexibility. Nature Reviews Neuroscience, 2012. 13(9): p. 636–650.

39. Thapar, A., et al., Depression in adolescence. Lancet, 2012. 379(9820): p. 1056–67.

40. Whitfield-Gabrieli, S. and J.M. Ford, Default mode network activity and connectivity in psychopathology. Annual review of clinical psychology, 2012. 8(1): p. 49–76.

41. Joormann, J. and I.H. Gotlib, Emotion regulation in depression: Relation to cognitive inhibition. Cognition and emotion, 2010. 24(2): p. 281–298.

42. Joormann, J. and W. Vanderlind, Emotion Regulation in Depression: The Role of Biased Cognition and Reduced Cognitive Control. Clinical Psychological Science, 2014. 2: p. 402–421.

43. Visted, E., et al., Emotion Regulation in Current and Remitted Depression: A Systematic Review and Meta-Analysis. Front Psychol, 2018. 9: p. 756.

44. Koenigs, M. and J. Grafman, The functional neuroanatomy of depression: distinct roles for ventromedial and dorsolateral prefrontal cortex. Behavioural brain research, 2009. 201(2): p. 239–243.

45. Hamilton, J.P., et al., Functional neuroimaging of major depressive disorder: a meta-analysis and new integration of baseline activation and neural response data. American Journal of Psychiatry, 2012. 169(7): p. 693–703.

46. Stange, J.P., L.B. Alloy, and D.M. Fresco, Inflexibility as a vulnerability to depression: A systematic qualitative review. Clinical psychology: Science and practice, 2017. 24(3): p. 245.

47. Drysdale, A.T., et al., Resting-state connectivity biomarkers define neurophysiological subtypes of depression. Nature medicine, 2017. 23(1): p. 28–38.

48. Pizzagalli, D.A., Depression, stress, and anhedonia: toward a synthesis and integrated model. Annual review of clinical psychology, 2014. 10(1): p. 393–423.

49. Hammen, C., Stress and depression. Annu. Rev. Clin. Psychol., 2005. 1: p. 293–319.

50. Slavich, G.M. and M.R. Irwin, From stress to inflammation and major depressive disorder: a social signal transduction theory of depression. Psychological bulletin, 2014. 140(3): p. 774.

51. Mayberg, H.S., et al., Deep brain stimulation for treatment-resistant depression. Neuron, 2005. 45(5): p. 651–660.

52. Fox, M.D., et al., Measuring and manipulating brain connectivity with resting state functional connectivity magnetic resonance imaging (fcMRI) and transcranial magnetic stimulation (TMS). Neuroimage, 2012. 62(4): p. 2232–2243.

53. Alexander, L.M., et al., An open resource for transdiagnostic research in pediatric mental health and learning disorders. Scientific Data, 2017. 4(1): p. 170181.

54. Kaufman, J., et al., Schedule for affective disorders and schizophrenia for school-age children-present and lifetime version (K-SADS-PL): initial reliability and validity data. Journal of the American Academy of Child & Adolescent Psychiatry, 1997. 36(7): p. 980–988.

55. Esteban, O., et al., fMRIPrep: a robust preprocessing pipeline for functional MRI. Nature methods, 2019. 16(1): p. 111–116.

56. Fischl, B., FreeSurfer. Neuroimage, 2012. 62(2): p. 774–781.

57. Avants, B.B., et al., Symmetric diffeomorphic image registration with cross-correlation: evaluating automated labeling of elderly and neurodegenerative brain. Medical image analysis, 2008. 12(1): p. 26–41.

58. Jenkinson, M., et al., Improved optimization for the robust and accurate linear registration and motion correction of brain images. Neuroimage, 2002. 17(2): p. 825–841.

59. Greve, D.N. and B. Fischl, Accurate and robust brain image alignment using boundary-based registration. Neuroimage, 2009. 48(1): p. 63–72.

60. Mehta, K., et al., XCP-D: A robust pipeline for the post-processing of fMRI data. Imaging Neuroscience, 2024. 2: p. imag-2-00257.

61. Satterthwaite, T.D., et al., An improved framework for confound regression and filtering for control of motion artifact in the preprocessing of resting-state functional connectivity data. Neuroimage, 2013. 64: p. 240–256.

62. Ciric, R., et al., Benchmarking of participant-level confound regression strategies for the control of motion artifact in studies of functional connectivity. Neuroimage, 2017. 154: p. 174–187.

63. Nastase, S.A., et al., Measuring shared responses across subjects using intersubject correlation. 2019, Oxford University Press. p. 667–685.

64. Hasson, U., et al., Brain-to-brain coupling: a mechanism for creating and sharing a social world. Trends in cognitive sciences, 2012. 16(2): p. 114–121.

65. Fisher, R.A., Frequency distribution of the values of the correlation coefficient in samples from an indefinitely large population. Biometrika, 1915. 10(4): p. 507–521.

66. Silver, N.C. and W.P. Dunlap, Averaging correlation coefficients: should Fisher’s z transformation be used? Journal of applied psychology, 1987. 72(1): p. 146.

67. Chen, G., et al., Untangling the relatedness among correlations, part I: Nonparametric approaches to inter-subject correlation analysis at the group level. NeuroImage, 2016. 142: p. 248–259.

68. Benjamini, Y. and Y. Hochberg, Controlling the false discovery rate: a practical and powerful approach to multiple testing. Journal of the Royal statistical society: series B (Methodological), 1995. 57(1): p. 289–300.

69. Smith, S.M. and T.E. Nichols, Threshold-free cluster enhancement: addressing problems of smoothing, threshold dependence and localisation in cluster inference. Neuroimage, 2009. 44(1): p. 83–98.

70. Chen, X., B. Lu, and C.G. Yan, Reproducibility of R-fMRI metrics on the impact of different strategies for multiple comparison correction and sample sizes. Human brain mapping, 2018. 39(1): p. 300–318.

71. Yan, C.-G., et al., DPABI: data processing & analysis for (resting-state) brain imaging. Neuroinformatics, 2016. 14(3): p. 339–351.

72. Winkler, A.M., et al., Faster permutation inference in brain imaging. Neuroimage, 2016. 141: p. 502–516.

73. Kim, D., et al., A new modular brain organization of the BOLD signal during natural vision. Cerebral Cortex, 2018. 28(9): p. 3065–3081.

74. Maris, E. and R. Oostenveld, Nonparametric statistical testing of EEG-and MEG-data. Journal of neuroscience methods, 2007. 164(1): p. 177–190.

75. Sassenhagen, J. and D. Draschkow, Cluster-based permutation tests of MEG/EEG data do not establish significance of effect latency or location. Psychophysiology, 2019. 56(6): p. e13335.

76. Gramfort, A., et al., MNE software for processing MEG and EEG data. neuroimage, 2014. 86: p. 446–460.

