## Supplementary Information for "Spatiotemporal unfolding of prefrontal neural response divergence in adolescent depression during naturalistic experience"

### Supplementary Methods

#### Movie clip annotation procedure

Cinematic annotations for each significant NPM episode identified in the dynamic neural polarization analysis were derived through a two-stage procedure. In the first stage, we manually reviewed the film footage corresponding to each episode and extracted the following information directly from the source material: (1) character dialogue; (2) background music and auditory atmosphere; and (3) salient visual changes, including key facial expressions, body language, and character actions. No inferential content was introduced at this stage; only observable elements explicitly present in the footage were recorded.

In the second stage, the manually extracted content was passed to a large language model (Gemini 3.1; Google DeepMind) using a structured prompt to derive standardized multi-dimensional narrative annotations. The prompt instructed the model to analyze each clip strictly on the basis of the provided material, without speculation or inference beyond what was explicitly documented. The full prompt used is reproduced below:

| Prompt (Gemini 3.1 thinking) |
|---|
| *Role*:<br>You are a professional film narrative and behavioral analyst. Your task is to extract the information specified below from raw film data I provide (sourced from *Despicable Me*), including character dialogue, visual presentation, and background music. All output must be in English and free from any pre-existing interpretive bias.<br><br>*Rules:*<br>● No speculation: extract only information explicitly present in the provided material.<br>● Skip missing items: if background music or specific dialogue is not mentioned in the source material, omit that field entirely without generating a placeholder.<br>● Be concise: each section should not exceed three lines.<br><br>*Output requirements (for each clip):*<br>**[Title]** Assign a brief descriptive title to facilitate reference.<br>**[Main Event]** Summarize the core narrative event of the clip.<br>**[Dialogue]** Extract the most salient dialogue. What do the characters say?<br>**[Details]** Extract important scene details, including character actions and facial expressions, character stance, and atmosphere (e.g., presence of background music, shifts in mood or tone).<br>**[Narrative Dimensions]** Characterize the clip along the following two dimensions:<br>- *Social Interaction:* Does the clip involve interpersonal evaluation, sense of belonging, social exclusion, or cooperation?<br>- *Emotional Valence*: Does the narrative convey a predominantly positive, negative, or neutral/ambivalent emotional tone?<br>**[Key Narrative Label]** Assign 1–2 descriptive labels to the clip (e.g., social evaluation, emotional resonance, action logic, self-reflection). |

### Supplementary Results

#### Within-group ISC exceeds between-group ISC in both polarized clusters across diagnostic groups

To confirm that the neural polarization identified in the dmPFC-rACC and left dlPFC clusters was not driven by a single diagnostic group, we conducted a confirmatory analysis examining whether within-group ISC exceeded between-group ISC independently in each group. For each cluster, voxel-wise within-group and between-group ISC values were averaged across all significant voxels to derive cluster-level ISC metrics. One-tailed paired $t$-tests were then conducted separately within the depression (MDD) and control (HC) groups to assess whether within-group ISC was significantly greater than between-group ISC. To further examine whether the magnitude of this within-versus-between difference was modulated by diagnostic group, an independent-samples $t$-test was applied to the difference scores ($ISC_{\text{within}} - ISC_{\text{between}}$) between the two groups (Supplementary Fig. S1).

In the dmPFC-rACC cluster, within-group ISC was significantly greater than between-group ISC in both the MDD group ($t(35) = 5.787, P < 0.001$) and the HC group ($t(35) = 4.975, P < 0.001$). The magnitude of the within-versus-between difference did not differ significantly between the two groups ($t(70) = 0.572, P = 0.569$), indicating that both groups contributed comparably to the observed neural polarization in this cluster.

In the left dlPFC cluster, within-group ISC also significantly exceeded between-group ISC in both the MDD group ($t(35) = 8.724, P < 0.001$) and the HC group ($t(35) = 2.262, P = 0.015$). However, the within-versus-between difference was significantly larger in the MDD group than in the HC group ($t(70) = 4.689, P < 0.001$), suggesting that while polarization was present in both groups, the internal neural coherence within the MDD group was substantially stronger in this cluster. Collectively, these results confirm that the neural polarization in both clusters reflects group-specific synchronization rather than a pattern driven by one diagnostic group alone.

#### Whole-brain representational similarity analysis

The primary neural polarization analyses treated the MDD–HC distinction as a categorical variable, identifying voxels where response similarity was greater within than between diagnostic groups. An alternative approach is to treat depressive symptom severity as a continuous measure and use representational similarity analysis (RSA) [1] to identify voxels where greater pairwise differences in symptom severity correspond to greater pairwise neural dissimilarity. We ran a whole-brain voxel-wise RSA to examine whether this continuous, graded relationship was detectable during film viewing.

The behavioral representational dissimilarity matrix (RDM) was constructed by computing the absolute pairwise difference in MFQ-SR total scores across all 2,556 unique participant pairs ($N = 72$). At each voxel, the neural RDM was defined as the pairwise neural dissimilarity, 1 − Pearson $r$, between the BOLD timecourses of every participant pair. A Pearson correlation was then computed between the neural RDM vector and the behavioral RDM vector at each voxel, with pairwise differences in age, sex, and mean framewise displacement included as covariates. Statistical inference used the same TFCE-corrected permutation procedure as the primary analysis (one-tailed, $P < 0.05$, 10,000 permutations; *Methods*). All voxel-wise correlation analyses and TFCE-based inference were implemented in the DPABI toolbox [2, 3].

The RSA yielded 11 significant clusters comprising 561 voxels in total (Supplementary Fig. S2). The two largest clusters were localized in the left lingual gyrus (cluster 1; 211 voxels; predominantly cLinG_L and rLinG_L; $r = 0.070 - 0.150$) and the right posterior superior temporal sulcus (cluster 3; 173 voxels; predominantly rpSTS_R, aSTS_R, and A37dl_R; $r = 0.068 - 0.144$). A third cluster (cluster 8; 80 voxels; predominantly A32p_L, A9m_L, and A32p_R; $r = 0.073 - 0.114$) fell within the medial prefrontal cortex and anterior cingulate cortex. The remaining eight clusters each comprised 27 voxels or fewer, distributed across temporal, parietal, and motor cortices (Supplementary Table S5). Spatial overlap between the RSA and primary neural polarization results was limited to 52 voxels, all falling within RSA cluster 8 and the dmPFC-rACC cluster (predominantly A32p, A8m, and A9m); no overlap was observed with the left dlPFC cluster.

The RSA and neural polarization analyses test different hypotheses. The RSA asks whether participants who differ more in symptom severity also differ more in their neural responses—a continuous relationship that treats all 2,556 participant pairs symmetrically and makes no assumption about diagnostic categories. The neural polarization analysis, by contrast, tests whether neural responses are more aligned within diagnostic groups than between them, a categorical formulation that does not require any linear mapping between symptom load and neural distance.

The two approaches also differ in how they pool information across observations. In the neural polarization analysis, each participant's polarization estimate averages pairwise ISC values across approximately 35 within-group and 36 between-group pairs before the group-level statistical test, substantially reducing measurement noise at each voxel. The RSA enters all 2,556 individual pairwise timecourse correlations as independent observations; individual-level pairwise correlations are considerably noisier than within-group means, which reduces effective sensitivity.

The RSA signal concentrated in the left lingual gyrus and right pSTS likely reflects continuous individual differences in visual and auditory-social processing that scale with symptom load but do not produce a categorical group split large enough to survive the neural polarization threshold. The absence of an RSA signal in the left dlPFC is consistent with the non-significant within-group symptom-severity correlations in the depression group (Supplementary Fig. S3), suggesting that the depression-related pattern in this cluster is better characterized as a discrete boundary than a graded continuum. The 52-voxel overlap between RSA cluster 8 and the dmPFC-rACC (A32p, A8m, A9m) indicates that medial prefrontal cortex carries both forms of depression-related neural variation, providing independent corroborating evidence for the primary dmPFC-rACC finding.

#### Cluster-level validation of dynamic neural polarization

The primary dynamic analysis quantified neural polarization using a voxel-wise approach, in which the ISPS-based polarization index was computed independently for each voxel and significant temporal windows were identified via MNE temporal cluster-based permutation testing; the summed polarization intensity across significant voxels at each time point yielded the NPM trajectory (see *Dynamic Neural Polarization Analysis* in the Methods). To evaluate the robustness of the identified temporal patterns, we conducted two complementary cluster-level analyses that aggregate neural signals prior to statistical testing.

In the first approach (*cluster-averaged BOLD*), the BOLD time series of all voxels within each cluster were spatially averaged to yield a single representative time series per cluster per participant. Instantaneous phase was then extracted via the Hilbert transform, pairwise ISPS was computed between participants, and the within-group minus between-group difference was derived at each time point, yielding a cluster-level dynamic polarization time series. Significant temporal windows were identified using the same MNE temporal cluster-based permutation procedure as the primary analysis (one-tailed, $P < 0.05$, 10,000 permutations; Supplementary Fig. S5).

In the second approach (*cluster-averaged ISPS*), the voxel-wise ISPS polarization values—computed identically to the primary analysis—were spatially averaged across all voxels within each cluster at each time point, producing a cluster-level mean phaseNDV time series per participant. The same temporal cluster correction was then applied (Supplementary Fig. S6).

Because the NPM approach aggregates voxels that individually achieve temporal significance, it can detect polarization driven by a spatial subset of the cluster at any given time point. Spatial averaging prior to testing, by contrast, mixes signal from both polarizing and non-polarizing voxels, reducing sensitivity to spatially focal effects. As expected, under these conditions, the significant temporal windows identified by both cluster-level approaches recovered a subset of the primary NPM episodes. For the cluster-averaged BOLD approach, significant episodes were identified in the dmPFC-rACC during TR 10–33 and TR 202–224, and in the left dlPFC during TR 5–23, TR 33–48, and TR 590–603. For the cluster-averaged ISPS approach, significant episodes were observed in the dmPFC-rACC during TR 195–234, and in the left dlPFC during TR 4–24, TR 31–51, and TR 584–604. Inspection of the full cluster-level trajectories further reveals that non-significant windows also exhibit peaks at time points corresponding to the primary NPM peaks, indicating that the underlying temporal structure of neural polarization is preserved across aggregation strategies even where individual peaks do not survive the significance threshold. The consistency in peak timing across all three analytical approaches supports the reliability of the dynamic polarization patterns reported in the main analysis.

#### Supplementary Figures

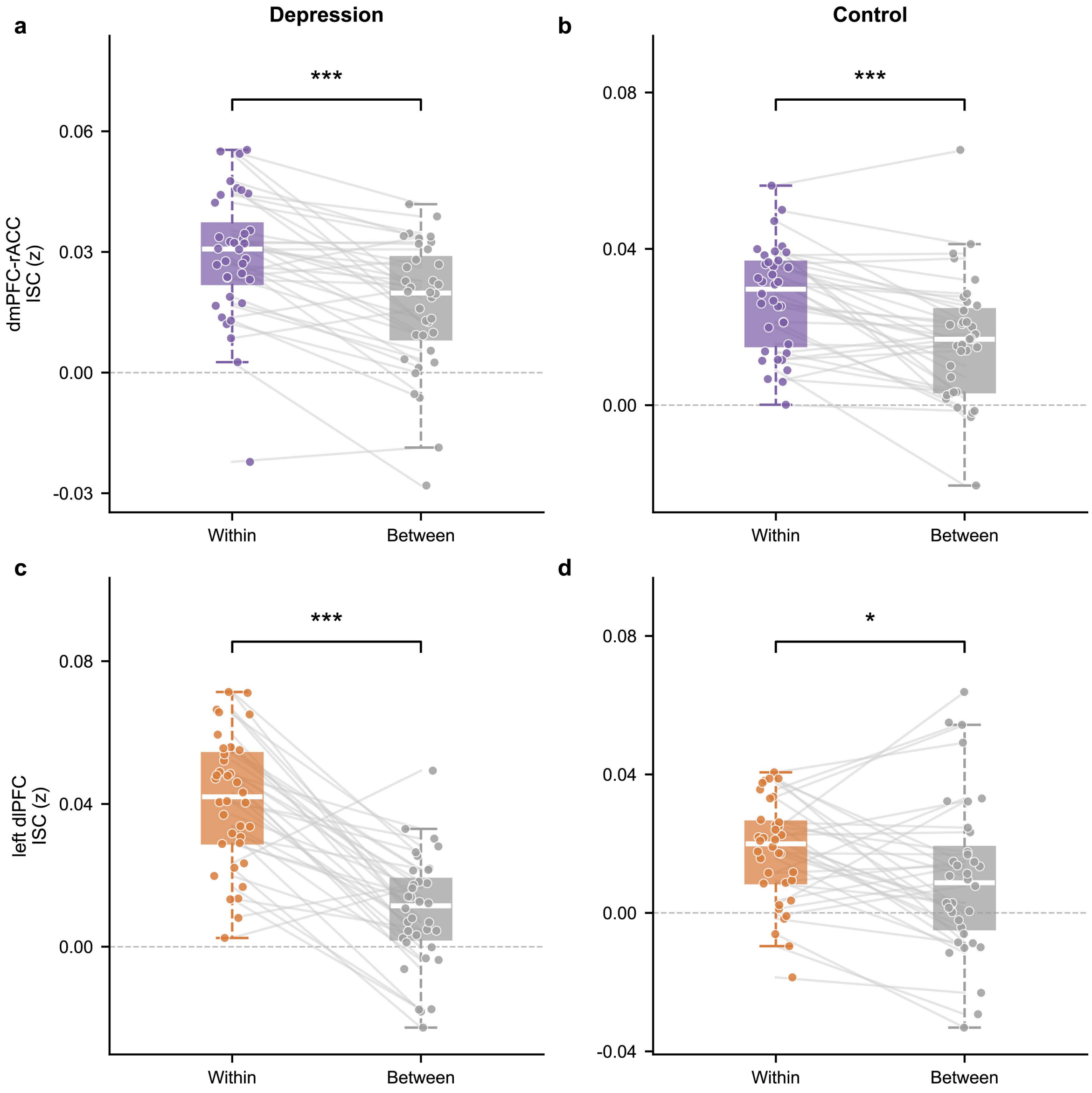


**Fig. S1. Within-group ISC exceeds between-group ISC in both diagnostic groups.** Paired comparisons of within-group and between-group inter-subject correlation (ISC) for the dmPFC-rACC (a, b) and left dlPFC (c, d) clusters, shown separately for the depression (n = 36; a, c) and control (n = 36; b, d) groups. Boxes indicate the median and interquartile range; lines connect paired values for each participant. Statistical significance was assessed using one-tailed paired t-tests (within > between). dmPFC-rACC: Depression, $t(35) = 5.79$, $P < 0.001$ (a); Control, $t(35) = 4.97$, $P < 0.001$ (b). left dlPFC: Depression, $t(35) = 8.72$, $P < 0.001$ (c); Control, $t(35) = 2.26$, $P = 0.015$ (d). ***uncorrected $P < 0.001$; *uncorrected $P < 0.05$.

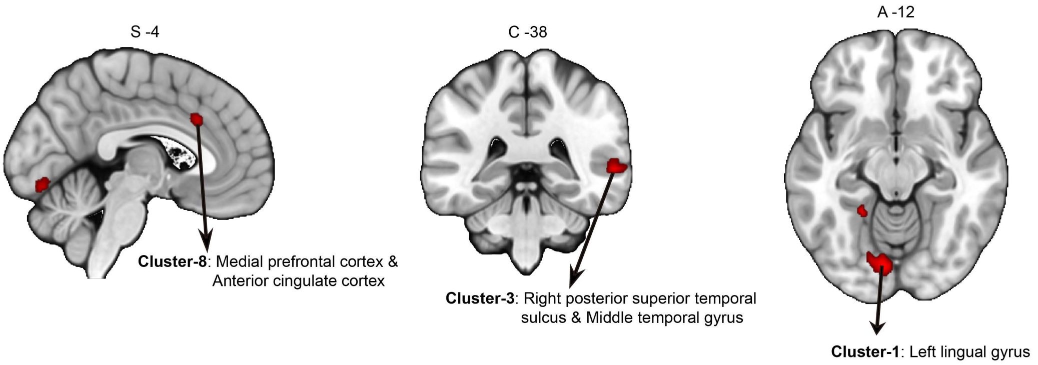


**Fig. S2. Whole-brain RSA results: voxels showing a significant positive association between pairwise neural dissimilarity and pairwise differences in depressive symptom severity.** Three representative sections were selected to highlight the three largest clusters. Left (sagittal, S = −4): Cluster-8, medial prefrontal cortex and anterior cingulate cortex (80 voxels; predominantly A32p, A9m, and A8m; r = 0.073–0.114). Middle (coronal, C = −38): Cluster-3, right posterior superior temporal sulcus and middle temporal gyrus (173 voxels; predominantly rpSTS_R and aSTS_R; r = 0.068–0.144). Right (axial, A = −12): Cluster-1, left lingual gyrus (211 voxels; predominantly cLinG_L; r = 0.070–0.150). Full cluster statistics and Brainnetome atlas parcellation for all clusters are provided in Supplementary Table S5.

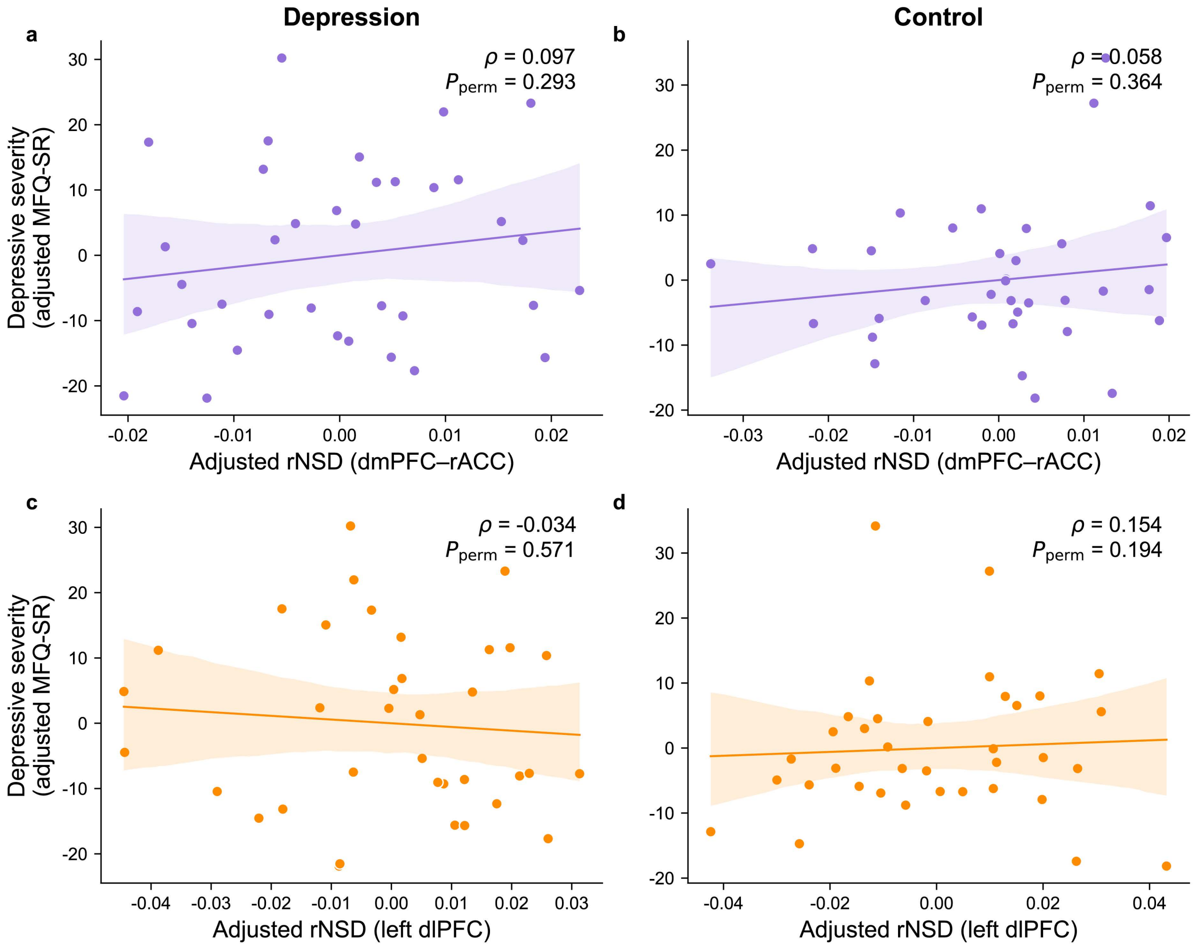


**Fig. S3. rNSD–symptom correlations are not significant within individual diagnostic groups.** Residual scatter plots showing the within-group association between rNSD scores and depressive symptom severity (MFQ-SR) for the dmPFC-rACC (a, b) and left dlPFC (c, d), computed separately within the depression (n = 36; a, c) and control (n = 36; b, d) groups. The rNSD index reflects each participant's neural similarity to the depression group relative to the control group (see *Methods*).Both axes reflect residuals after regressing out age, sex, and mean framewise displacement; shaded bands indicate 95% confidence intervals. Statistical associations were assessed using partial Spearman's correlation controlling for age, sex, and mean FD; *P* values were derived from nonparametric permutation tests (10,000 iterations). No significant association was observed in either group for either ROI (dmPFC-rACC: Depression, $\rho = 0.097$, $P_{perm} = 0.293$; Control, $\rho = 0.058$, $P_{perm} = 0.364$; left dlPFC: Depression, $\rho = -0.034$,$P_{perm} = 0.571$; Control, $\rho = 0.154$, $P_{perm} = 0.194$). rNSD, relative neural similarity to depression; MFQ-SR, Moods and Feelings Questionnaire (self-report).

**Clip3: Proposal to Delay the Heist**

[**Main Event**] Gru hesitantly suggests moving the heist date to Dr. Nefario, who suspects the girls' dance recital is the cause.

[**Details**] Gru tidies toys on the floor with his back turned to the Doctor while being followed and questioned.

[**Social Interaction**] Subtle interpersonal negotiation and evasion with a collaborator.

[**Emotional Valence**] ambivalent/neutral; characterized by hesitation.

[**Narrative Label**] interpersonal negotiation.

*Gru: "Uh… About that, I was thinking that maybe we could move the date of the heist."*
*Dr. Nefario: "Please tell me this is not as a result of the girls' dance recital, is it?"*

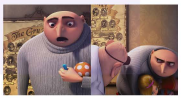

**Clip5: Minions' Absurdist Humor**

[**Main Event**] Following a heavy emotional scene, the focus shifts abruptly to Minions engaging in silly behavior by photocopying their bottoms and laughing.

[**Details**] A sudden transition from a dark to a bright visual setting; Minions are shown sharing a playful moment.

[**Social Interaction**] Peer-level shared amusement.

[**Emotional Valence**] positive; sudden mood shift from the previous tension.

[**Narrative Label**] Humorous Relief; Mood Shift.

*Minions: "Butt" followed by continuous, high-pitched laughter.*

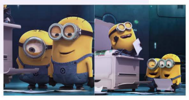

**Clip6: Hostile Confrontation**

[**Main Event**] Miss Hattie arrives to take the girls back and physically strikes Gru with a dictionary.

[**Details**] Miss Hattie strikes Gru across the face with a heavy dictionary. Gru gasps and appears shocked as Dr. Nefario emerges behind him.

[**Social Interaction**] Hostile social encounter involving physical aggression.

[**Emotional Valence**] negative; shock and hostility.

[**Narrative Label**] physical aggression; social conflict.

*Miss Hattie: "I received a call that you wanted to return them... I didn't like what you said."*

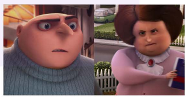

**Clip9: The Ticket Distraction (same as Clip13)**

[**Main Event**] Gru's focused mission preparation is interrupted by a Minion delivering a dance recital ticket.

[**Details**] Gru stands on a mission lift, transitioning from a determined, heroic pose to an expression of annoyance.

[**Social Interaction**] Personal social reminder intrudes upon a high-stakes professional role.

[**Emotional Valence**] negative; characterized by annoyance.

[**Narrative Label**] cognitive interference; task switching.

*Gru: "What is this for?"*

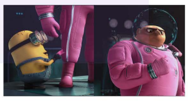

**Fig. S4. Additional cinematic episodes driving significant neural polarization in the dmPFC-rACC.** Four additional temporal segments producing significant neural polarization magnitude (NPM) in the dmPFC-rACC hub, supplementing the primary episodes described in Fig. 5. Panel format follows Fig. 5: each entry provides the primary cinematic event, social interaction type, emotional valence, narrative label, and a representative still frame.

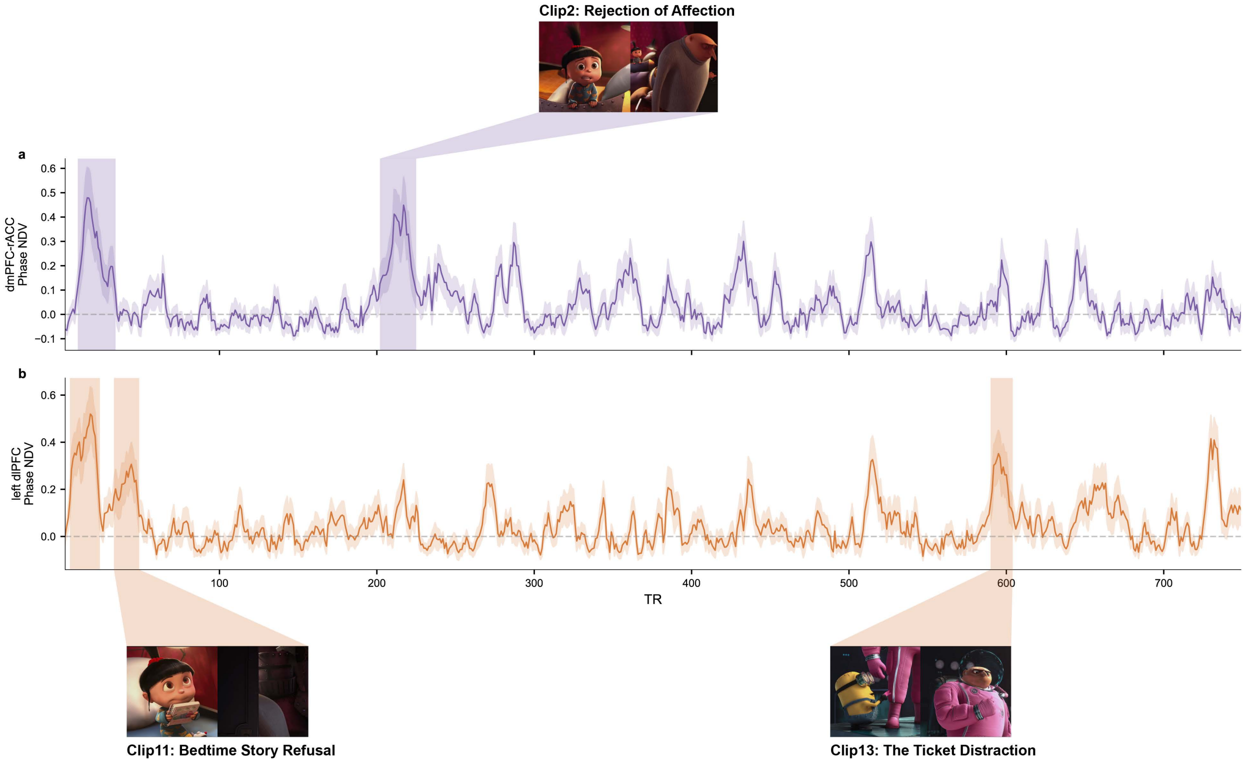


**Fig. S5. Dynamic neural polarization trajectories derived from cluster-averaged BOLD signals.** Each panel displays the group-mean phase-based neural divergence (phase NDV) time series across the full scan duration (TR 2–749, n = 72), computed from the spatially averaged BOLD signal within each cluster. For each participant, the BOLD time series of all voxels within the cluster were averaged to yield a single representative time series, from which instantaneous phase was extracted via the Hilbert transform. Phase NDV at each time point was defined as the within-group minus between-group inter-subject phase synchrony (ISPS), Fisher z-transformed and averaged across participant pairs. Shaded bands indicate significant temporal clusters identified by nonparametric temporal cluster-based permutation testing (one-tailed, cluster-forming threshold $P < 0.05$, 10,000 permutations). **a** dmPFC-rACC (purple); significant episodes: TR 10–33, TR 202–224. **b** Left dlPFC (orange); significant episodes: TR 5–23, TR 33–48, TR 590–603. Shaded ribbons around the mean line indicate ±1 SEM across participants. Representative still frames and clip labels mark the cinematic episodes corresponding to primary NPM peaks; detailed narrative annotations for each episode are provided in Fig. 5 and the main text.

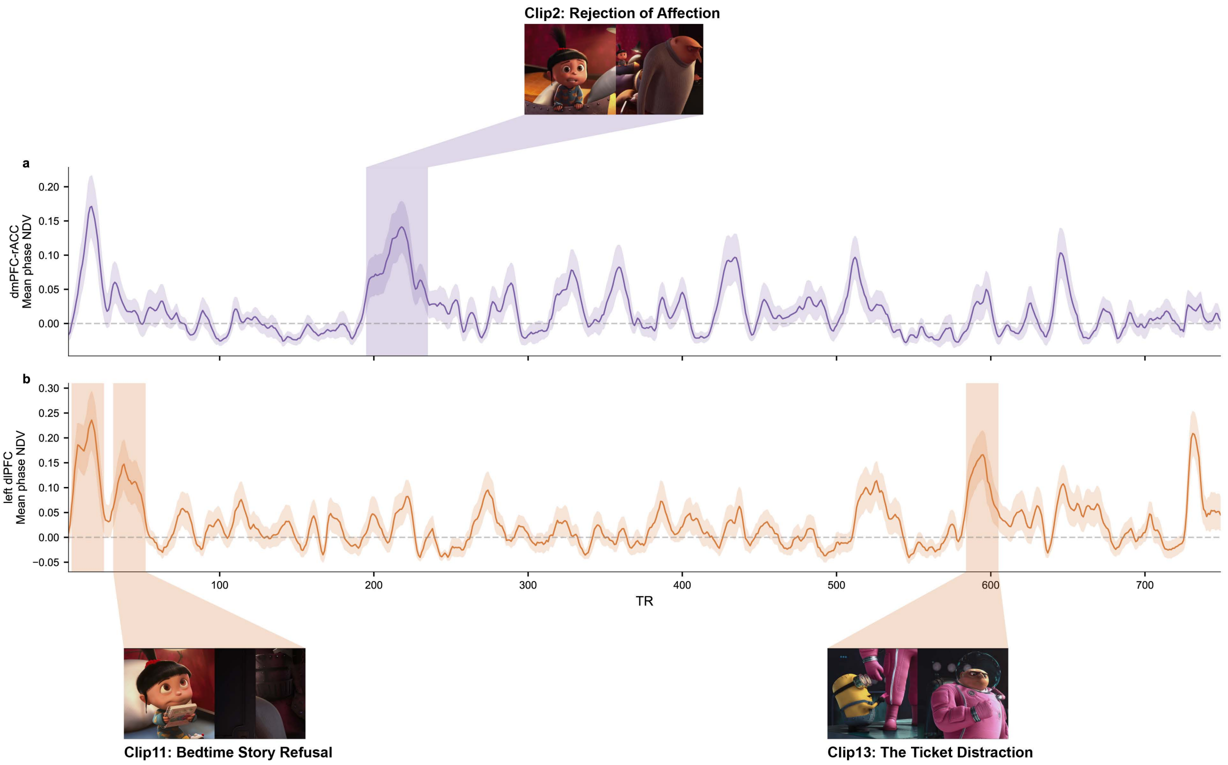


**Fig. S6. Dynamic neural polarization trajectories derived from cluster-averaged ISPS values.** Each panel displays the group-mean phase NDV time series across the full scan duration (TR 2–749, n = 72), computed by spatially averaging voxel-wise ISPS polarization values across all voxels within each cluster at each time point. Voxel-wise phase NDV was computed identically to the primary analysis (within-group minus between-group ISPS, Fisher z-transformed). Shaded bands indicate significant temporal clusters identified by nonparametric temporal cluster-based permutation testing (one-tailed, cluster-forming threshold $P < 0.05$, 10,000 permutations). **a** dmPFC-rACC (purple); significant episode: TR 195–234. **b** Left dlPFC (orange); significant episodes: TR 4–24, TR 31–51, TR 584–604. Shaded ribbons around the mean line indicate ±1 SEM across participants. Representative still frames and clip labels mark the cinematic episodes corresponding to primary NPM peaks; detailed narrative annotations for each episode are provided in Fig. 5 and the main text.

#### Supplementary Tables

**Table S1. Cluster statistics and Brainnetome atlas parcellation of the dmPFC-rACC neural polarization cluster.**

| Cluster level | | | | |
|---|---|---|---|---|
| **Cluster label** | **Cluster size** | **MNI (X,Y,Z) (peak)** | **Fisher's $z$-value (peak)** | **$t$-value (peak)** |
| dmPFC-rACC | 618 | (2, 32, 24) | 0.020 | 5.211 |

| Voxel level | | | | | | |
|---|---|---|---|---|---|---|
| **Atlas ID** | **Name** | **$N_{voxel}$** | **Percent** | **Mean $t$ (range)** | **Anatomical and modified Cyto-architectonic descriptions** | **Gyrus** |
| 1 | A8m_L | 32 | 5.178 | 3.367 (2.843–4.814) | A8m, medial area 8 | SFG |
| 2 | A8m_R | 3 | 0.485 | 3.157 (2.914–3.347) | | SFG |
| 11 | A9m_L | 78 | 12.621 | 3.524 (2.839–4.446) | A9m,medial area 9 | SFG |
| 12 | A9m_R | 50 | 8.091 | 3.346 (2.852–4.245) | | SFG |
| 13 | A10m_L | 4 | 0.647 | 3.079 (2.848–3.313) | A10m, medial area 10 | SFG |
| 14 | A10m_R | 12 | 1.942 | 3.847 (3.069–5.163) | | SFG |
| 177 | A24rv_L | 5 | 0.809 | 3.185 (3.020–3.431) | A24rv, rostroventral area 24 | CG |
| 178 | A24rv_R | 12 | 1.942 | 3.289 (2.917–3.702) | | CG |
| 179 | A32p_L | 123 | 19.903 | 3.627 (2.865–4.997) | A32p, pregenual area 32 | CG |
| 180 | A32p_R | 159 | 25.728 | 3.64 (2.854–5.211) | | CG |
| 183 | A24cd_L | 38 | 6.149 | 3.536 (2.904–4.684) | A24cd, caudodorsal area 24 | CG |
| 184 | A24cd_R | 36 | 5.825 | 3.545 (2.970–4.473) | | CG |
| 187 | A32sg_L | 17 | 2.751 | 3.094 (2.843–3.583) | A32sg, subgenual area 32 | CG |
| 188 | A32sg_R | 49 | 7.929 | 3.654 (2.855–5.041) | | CG |

The upper section (cluster level) summarizes the full voxel cluster identified by the static neural polarization analysis: cluster label, size (voxels), MNI peak coordinates (x, y, z; mm), peak Fisher's z-transformed ISC difference, and peak voxel-wise t-statistic. The lower section (voxel level) details the distribution of voxels within the cluster across Brainnetome atlas parcels. For each parcel, the Atlas ID and anatomical name follow the Brainnetome atlas nomenclature [4]. $N_{voxel}$ gives the number of cluster voxels falling within each parcel; % of cluster is the corresponding proportion of the 618-voxel cluster total. Mean $t$ (range) reports the mean voxel-wise t-statistic across all voxels in that parcel, with the minimum and maximum values shown in parentheses. Gyrus indicates the macro-anatomical label. The cluster label, dmPFC-rACC, reflects the predominant contribution of A9m (medial area 9, dorsomedial prefrontal cortex; 20.71%) and A32p (pregenual area 32, rostral anterior cingulate cortex; 45.63%), together accounting for the majority of cluster voxels. MNI, Montreal Neurological Institute; SFG, superior frontal gyrus; CG, cingulate gyrus.

**Table S2. Cluster statistics and Brainnetome atlas parcellation of the left dlPFC neural polarization cluster.**

**Cluster level**

| Cluster label | Cluster size | MNI (X,Y,Z) (peak) | Fisher's $z$-value (peak) | $t$-value (peak) |
|---|---|---|---|---|
| Left dlPFC | 142 | (-24,52,30) | 0.030 | 6.853 |

**Voxel level**

| Atlas ID | Name | $N_{voxel}$ | Percent | Mean $t$ (range) | Anatomical and modified Cyto-architectonic descriptions | Gyrus |
|---|---|---|---|---|---|---|
| 5 | A9l_L | 1 | 0.704 | 3.842 (3.842–3.842) | A9l, lateral area 9 | SFG |
| 15 | A9/46d_L | 141 | 99.296 | 4.803 (3.688–6.853) | A9/46d, dorsal area 9/46 | MFG |

The upper section (cluster level) summarizes the left dlPFC voxel cluster identified by the static neural polarization analysis: cluster label, size (voxels), MNI peak coordinates (x, y, z; mm), peak Fisher's z-transformed ISC difference, and peak voxel-wise t-statistic. The lower section (voxel level) details the distribution of voxels within the cluster across Brainnetome atlas parcels. For each parcel, the Atlas ID and anatomical name follow the Brainnetome atlas. $N_{voxel}$ gives the number of cluster voxels falling within each parcel; % of cluster is the corresponding proportion of the 142-voxel cluster total. Mean $t$ (range) reports the mean voxel-wise t-statistic across all voxels in that parcel, with the minimum and maximum values shown in parentheses. Gyrus indicates the macro-anatomical label. The cluster label, left dlPFC, reflects the near-exclusive contribution of A9/46d_L (dorsal area 9/46; 99.3%), a cytoarchitectonically defined region within the left middle frontal gyrus corresponding to the dorsolateral prefrontal cortex. MNI, Montreal Neurological Institute; SFG, superior frontal gyrus; MFG, middle frontal gyrus.

**Table S3. Meta-analytic functional decoding term rankings for the dmPFC-rACC cluster.**

| Term | probReverse | zReverse | pReverse |
|---|---|---|---|
| painful | 0.7192 | 8.027 | 1.000E-15 |
| insula anterior | 0.7170 | 6.766 | 1.325E-11 |
| noxious | 0.7117 | 5.259 | 1.451E-07 |
| nociceptive | 0.7117 | 4.772 | 1.827E-06 |
| anterior cingulate | 0.7113 | 19.578 | 2.361E-85 |
| acc | 0.7054 | 11.770 | 5.574E-32 |
| signal task | 0.7048 | 5.052 | 4.368E-07 |
| anterior insular | 0.7017 | 4.873 | 1.099E-06 |
| cortex acc | 0.6990 | 9.771 | 1.501E-22 |
| compensation | 0.6913 | 4.492 | 7.063E-06 |
| dorsal anterior | 0.6912 | 9.172 | 4.634E-20 |
| impulsivity | 0.6900 | 5.085 | 3.674E-07 |
| stop signal | 0.6892 | 4.600 | 4.219E-06 |
| cingulate | 0.6878 | 18.297 | 8.741E-75 |
| pain | 0.6756 | 8.926 | 4.403E-19 |
| salience network | 0.6727 | 4.490 | 7.121E-06 |
| rostral anterior | 0.6726 | 4.063 | 4.837E-05 |
| dacc | 0.6710 | 4.997 | 5.838E-07 |
| sn | 0.6672 | 3.381 | 7.210E-04 |
| cingulate cortex | 0.6652 | 13.700 | 1.012E-42 |
| cortex dacc | 0.6640 | 4.107 | 4.003E-05 |
| anterior | 0.6612 | 15.504 | 3.236E-54 |
| autonomic | 0.6601 | 3.870 | 1.089E-04 |
| matrix | 0.6589 | 3.272 | 1.066E-03 |
| cortex frontal | 0.6585 | 3.106 | 1.898E-03 |
| whilst | 0.6574 | 3.528 | 4.194E-04 |
| conflict | 0.6562 | 6.202 | 5.565E-10 |
| serves | 0.6559 | 3.128 | 1.759E-03 |
| risk taking | 0.6559 | 3.128 | 1.759E-03 |
| cortex insula | 0.6558 | 4.059 | 4.935E-05 |
| impulsive | 0.6556 | 3.042 | 2.348E-03 |
| rostral | 0.6532 | 6.346 | 2.210E-10 |
| involved cognitive | 0.6522 | 2.890 | 3.855E-03 |
| periaqueductal | 0.6498 | 2.918 | 3.522E-03 |
| reappraisal | 0.6484 | 3.035 | 2.404E-03 |
| anterior insula | 0.6463 | 7.791 | 6.637E-15 |
| cingulate cortices | 0.6462 | 3.617 | 2.985E-04 |
| stop | 0.6451 | 3.808 | 1.400E-04 |
| distress | 0.6451 | 2.889 | 3.867E-03 |
| skin conductance | 0.6433 | 2.921 | 3.489E-03 |

| | | | |
|---|---|---|---|
| competing | 0.6427 | 3.524 | 4.258E-04 |
| neurobiology | 0.6427 | 3.040 | 2.366E-03 |
| frontal operculum | 0.6421 | 3.155 | 1.607E-03 |
| error | 0.6412 | 6.250 | 4.105E-10 |
| networks involved | 0.6409 | 3.186 | 1.445E-03 |
| cortex thalamus | 0.6397 | 2.774 | 5.539E-03 |
| lateral medial | 0.6390 | 2.544 | 1.096E-02 |
| response inhibition | 0.6388 | 4.275 | 1.914E-05 |
| cortex anterior | 0.6384 | 5.334 | 9.629E-08 |
| conductance | 0.6383 | 2.810 | 4.959E-03 |
| stroop task | 0.6359 | 3.342 | 8.318E-04 |
| preferences | 0.6358 | 2.755 | 5.868E-03 |
| cingulate gyrus | 0.6353 | 5.172 | 2.317E-07 |
| regard | 0.6351 | 3.265 | 1.094E-03 |
| conflicting | 0.6337 | 2.954 | 3.140E-03 |
| losses | 0.6333 | 2.701 | 6.911E-03 |
| primary secondary | 0.6333 | 2.701 | 6.911E-03 |
| prefrontal cortical | 0.6333 | 2.701 | 6.911E-03 |
| network connectivity | 0.6329 | 3.362 | 7.740E-04 |
| learning task | 0.6320 | 3.025 | 2.484E-03 |
| motivational | 0.6320 | 3.757 | 1.719E-04 |
| 32 | 0.6292 | 3.959 | 7.522E-05 |
| control processes | 0.6292 | 3.311 | 9.300E-04 |
| somatosensory cortices | 0.6291 | 2.729 | 6.360E-03 |
| engages | 0.6288 | 3.051 | 2.278E-03 |
| middle cingulate | 0.6287 | 2.409 | 1.598E-02 |
| zone | 0.6279 | 2.258 | 2.397E-02 |
| producing | 0.6277 | 2.454 | 1.411E-02 |
| prediction error | 0.6268 | 2.499 | 1.246E-02 |
| empathic | 0.6268 | 2.499 | 1.246E-02 |

The table lists the top 70 terms associated with the dmPFC-rACC spatial mask, derived from meta-analytic functional decoding using the Neurosynth $\chi^2$ method. Terms are ranked in descending order of reverse-inference probability (probReverse). The top 20 functional-related terms from this ranking are displayed in the word cloud adjacent to Fig. 3a, with font size scaled by zReverse. probReverse is the posterior probability that a given term appears in a study, conditioned on activation within the specified brain region (P(term | activation)). zReverse is the standardized effect size of this reverse-inference association, correcting for the base rate of each term across the Neurosynth database. pReverse is the corresponding one-tailed uncorrected p-value.

**Table S4. Meta-analytic functional decoding term rankings for the left dlPFC cluster.**

| Term | probReverse | zReverse | pReverse |
|---|---|---|---|
| children adolescents | 0.8620 | 5.089 | 3.599E-07 |
| eating | 0.8484 | 5.073 | 3.926E-07 |
| nociceptive | 0.8469 | 4.277 | 1.898E-05 |
| maximal | 0.8469 | 4.277 | 1.898E-05 |
| midline | 0.8307 | 5.705 | 1.161E-08 |
| cortex insula | 0.8237 | 4.729 | 2.251E-06 |
| young older | 0.8169 | 3.263 | 1.103E-03 |
| skin conductance | 0.8159 | 3.618 | 2.964E-04 |
| dorsolateral pfc | 0.8126 | 3.556 | 3.766E-04 |
| conductance | 0.8126 | 3.556 | 3.766E-04 |
| analyzing | 0.8087 | 3.124 | 1.783E-03 |
| repeat | 0.8087 | 3.124 | 1.783E-03 |
| gyrus anterior | 0.8023 | 4.225 | 2.389E-05 |
| varies | 0.7986 | 2.964 | 3.040E-03 |
| serves | 0.7966 | 2.933 | 3.356E-03 |
| validity | 0.7933 | 3.215 | 1.305E-03 |
| neutral pictures | 0.7926 | 2.873 | 4.061E-03 |
| network connectivity | 0.7913 | 3.744 | 1.813E-04 |
| enhancing | 0.7907 | 2.844 | 4.452E-03 |
| time courses | 0.7907 | 2.844 | 4.452E-03 |
| biased | 0.7886 | 3.137 | 1.704E-03 |
| 41 | 0.7868 | 2.787 | 5.317E-03 |
| drugs | 0.7868 | 2.787 | 5.317E-03 |
| cortex thalamus | 0.7848 | 2.759 | 5.793E-03 |
| noxious | 0.7829 | 2.732 | 6.299E-03 |
| conditioning | 0.7804 | 3.536 | 4.061E-04 |
| behavioral responses | 0.7772 | 2.652 | 8.011E-03 |
| hub | 0.7772 | 2.652 | 8.011E-03 |
| skin | 0.7760 | 3.458 | 5.449E-04 |
| connectivity networks | 0.7749 | 2.921 | 3.493E-03 |
| erp | 0.7736 | 3.169 | 1.530E-03 |
| indexed | 0.7734 | 2.600 | 9.322E-03 |
| courses | 0.7715 | 2.575 | 1.003E-02 |
| causality | 0.7697 | 2.550 | 1.078E-02 |
| emergence | 0.7690 | 2.206 | 2.736E-02 |
| 39 | 0.7689 | 2.831 | 4.639E-03 |
| food | 0.7686 | 3.326 | 8.811E-04 |
| high functioning | 0.7665 | 2.178 | 2.941E-02 |
| cortex dorsolateral | 0.7665 | 2.178 | 2.941E-02 |
| overlapped | 0.7645 | 2.766 | 5.670E-03 |

| | | | |
|---|---|---|---|
| economic | 0.7640 | 2.150 | 3.155E-02 |
| evolution | 0.7640 | 2.150 | 3.155E-02 |
| zone | 0.7640 | 2.150 | 3.155E-02 |
| imagine | 0.7640 | 2.150 | 3.155E-02 |
| cortex middle | 0.7623 | 2.453 | 1.416E-02 |
| salience network | 0.7615 | 2.724 | 6.447E-03 |
| choose | 0.7592 | 2.096 | 3.611E-02 |
| externally | 0.7587 | 2.407 | 1.610E-02 |
| amygdala anterior | 0.7568 | 2.069 | 3.853E-02 |
| sum | 0.7568 | 2.069 | 3.853E-02 |
| thinking | 0.7565 | 2.897 | 3.765E-03 |
| mean age | 0.7562 | 3.115 | 1.837E-03 |
| aversive | 0.7555 | 3.499 | 4.671E-04 |
| ventromedial | 0.7545 | 4.547 | 5.446E-06 |
| sn | 0.7544 | 2.043 | 4.105E-02 |
| retention | 0.7544 | 2.043 | 4.105E-02 |
| loop | 0.7544 | 2.043 | 4.105E-02 |
| consequences | 0.7543 | 3.289 | 1.006E-03 |
| refers | 0.7533 | 2.339 | 1.934E-02 |
| effectively | 0.7533 | 2.339 | 1.934E-02 |
| spectrum | 0.7530 | 3.629 | 2.844E-04 |
| somatosensory cortices | 0.7515 | 2.317 | 2.051E-02 |
| considering | 0.7515 | 2.317 | 2.051E-02 |
| images acquired | 0.7496 | 1.992 | 4.636E-02 |
| parieto occipital | 0.7496 | 1.992 | 4.636E-02 |
| modulatory | 0.7487 | 2.544 | 1.094E-02 |
| personal | 0.7481 | 2.984 | 2.843E-03 |
| joint | 0.7480 | 2.274 | 2.299E-02 |
| 20 healthy | 0.7480 | 2.274 | 2.299E-02 |
| granger | 0.7473 | 1.967 | 4.916E-02 |

The table lists the top 70 terms associated with the left dlPFC spatial mask, derived from meta-analytic functional decoding using the Neurosynth $\chi^2$ method. Terms are ranked in descending order of reverse-inference probability (probReverse). The top 20 functional-related terms from this ranking are displayed in the word cloud adjacent to Fig. 3b, with font size scaled by zReverse. probReverse is the posterior probability that a given term appears in a study, conditioned on activation within the specified brain region (P(term | activation)). zReverse is the standardized effect size of this reverse-inference association, correcting for the base rate of each term across the Neurosynth database. pReverse is the corresponding one-tailed uncorrected p-value.

**Table S5. Cluster statistics and Brainnetome atlas parcellation for the whole-brain representational similarity analysis (RSA).**

| Cluster level | | | | |
|---|---|---|---|---|
| **Cluster label** | **Description** | **Size** | **MNI (X,Y,Z) (peak)** | **$r$-value (peak)** |
| Cluster-1 | *Left lingual gyrus* | 211 | (-8, -80, -12) | 0.1497 |
| Cluster-2 | *Left fusiform & Parahippocampal gyrus* | 27 | (-22, -46, -14) | 0.1500 |
| Cluster-3 | *Right posterior superior temporal sulcus & Middle temporal gyrus* | 173 | (60, -38, 2) | 0.1437 |
| Cluster-4 | *Right middle temporal gyrus* | 10 | (54, -58, 2) | 0.1130 |
| Cluster-5 | *Left primary motor cortex (tongue/larynx representation)* | 8 | (-60, 4, 6) | 0.1193 |
| Cluster-6 | *Left occipital polar cortex* | 12 | (-8, -102, 12) | 0.1423 |
| Cluster-7 | *Right caudal area (A39c)* | 4 | (40, -58, 14) | 0.1293 |
| Cluster-8 | *Medial prefrontal cortex & Anterior cingulate cortex* | 80 | (-4, 22, 32) | 0.11352 |
| Cluster-9 | *Right superior parietal lobule* | 27 | (28, -60, 64) | 0.1344 |
| Cluster-10 | *Right superior parietal lobule* | 7 | (42, -46, 64) | 0.1579 |
| Cluster-11 | *Right superior parietal lobule* | 2 | (40, -48, 66) | 0.1646 |

| Voxel level (Cluster-1) | | | | | | |
|---|---|---|---|---|---|---|
| **Atlas ID** | **Name** | $N_{voxel}$ | **Percent** | **Mean $r$ (range)** | **Anatomical and modified Cyto-architectonic descriptions** | **Gyrus** |
| 189 | cLinG_L | 189 | 89.573 | 0.1018 (0.0698–0.1497) | *cLinG, caudal lingual gyrus* | *MVOcC* |
| 195 | rLinG_L | 22 | 10.427 | 0.0835 (0.0722–0.1011) | *rLinG, rostral lingual gyrus* | *MVOcC* |

| Voxel level (Cluster-2) | | | | | | |
|---|---|---|---|---|---|---|
| **Atlas ID** | **Name** | $N_{voxel}$ | **Percent** | **Mean $r$ (range)** | **Anatomical and modified Cyto-architectonic descriptions** | **Gyrus** |
| 103 | A20rv_L | 2 | 7.407 | 0.1080 (0.1010–0.1150) | *A20rv, rostroventral area 20* | *FuG* |
| 105 | A37mv_L | 14 | 51.852 | 0.1159 (0.0957–0.1501) | *A37mv, medioventral area37* | *FuG* |
| 119 | TH_L | 6 | 22.222 | 0.1063 (0.0931–0.1298) | *TH, area TH (medial PPHC)* | *PhG* |
| 195 | rLinG_L | 5 | 18.519 | 0.1125 (0.0982–0.1400) | *rLinG, rostral lingual gyrus* | *MVOcC* |

| Voxel level (Cluster-3) | | | | | | |
|---|---|---|---|---|---|---|
| **Atlas ID** | **Name** | $N_{voxel}$ | **Percent** | **Mean $r$ (range)** | **Anatomical and modified Cyto-architectonic descriptions** | **Gyrus** |
| 80 | A22r_R | 4 | 2.312 | 0.0794 (0.0763–0.0825) | *A22r, rostral area 22* | *STG* |
| 82 | A21c_R | 1 | 0.578 | 0.0687 (0.0687–0.0687) | *A21c, caudal area 21* | *MTG* |
| 86 | A37dl_R | 30 | 17.341 | 0.0790 (0.0683–0.0927) | *A37dl, dorsolateral area37* | *MTG* |
| 88 | aSTS_R | 51 | 29.48 | 0.0811 (0.0684–0.1059) | *A37dl, dorsolateral area37* | *MTG* |
| 122 | rpSTS_R | 75 | 43.353 | 0.0978 (0.0684–0.1437) | *rpSTS, rostroposterior superior temporal sulcus* | *pSTS* |
| 124 | cpSTS_R | 12 | 6.936 | 0.0777 (0.0693–0.0889) | *cpSTS, caudoposterior superior temporal sulcus* | *pSTS* |

| Atlas ID | Name | $N_{voxel}$ | Percent | Mean $r$ (range) | Anatomical and modified Cyto-architectonic descriptions | Gyrus |
|---|---|---|---|---|---|---|
| **Voxel level (Cluster-8)** | | | | | | |
| 1 | A8m_L | 11 | 13.75 | 0.0802 (0.0732–0.0886) | *A8m, medial area 8* | *SFG* |
| 2 | A8m_R | 8 | 10 | 0.0804 (0.0739–0.0930) | *A8m, medial area 8* | *SFG* |
| 11 | A9m_L | 12 | 15 | 0.0854 (0.0733–0.0974) | *A9m,medial area 9* | *SFG* |
| 179 | A32p_L | 25 | 31.25 | 0.0921 (0.0738–0.1135) | *A32p, pregenual area 32* | *CG* |
| 180 | A32p_R | 12 | 15 | 0.0795 (0.0756–0.0865) | *A32p, pregenual area 32* | *CG* |
| 183 | A24cd_L | 8 | 10 | 0.0872 (0.0796–0.1077) | *A24cd, caudodorsal area 24* | *CG* |
| 184 | A24cd_R | 4 | 5 | 0.0772 (0.0739–0.0845) | *A24cd, caudodorsal area 24* | *CG* |
| **Voxel level (Cluster-9)** | | | | | | |
| **Atlas ID** | **Name** | $N_{voxel}$ | **Percent** | **Mean $r$ (range)** | **Anatomical and modified Cyto-architectonic descriptions** | **Gyrus** |
| 126 | A7r_R | 6 | 22.222 | 0.1080 (0.0948–0.1344) | *A7r, rostral area 7* | *SPL* |
| 134 | A7ip_R | 21 | 77.778 | 0.1008 (0.0902–0.1199) | *A7ip, intraparietal area 7(hIP3)* | *SPL* |

The upper section (cluster level) summarizes all significant clusters identified in the whole-brain RSA, in which pairwise neural dissimilarity ($1 - \text{Pearson } r$) at each voxel was positively correlated with pairwise differences in MFQ-SR total scores ($N = 72$; 2,556 unique pairs), controlling for pairwise differences in age, sex, and mean framewise displacement. For each cluster, the table lists the cluster label, anatomical description, size (voxels), MNI peak coordinates (x, y, z; mm) and peak Pearson $r$-value. All clusters survived TFCE-corrected $P < 0.05$ (one-tailed, 10,000 permutations), reflecting a significant positive association between pairwise neural dissimilarity and pairwise differences in depressive symptom severity. The lower sections (voxel level) detail the Brainnetome atlas parcellation for each cluster exceeding 20 voxels; clusters with 20 or fewer voxels are listed in the cluster-level summary only. For each parcel, the Atlas ID and anatomical name follow the Brainnetome atlas nomenclature. $N_{voxel}$ gives the number of cluster voxels falling within each parcel; % of cluster is the corresponding proportion of the cluster total. Mean $r$ (range) reports the mean Pearson $r$-value across all voxels in that parcel, with the minimum and maximum values in parentheses. Gyrus and Lobe indicate macro-anatomical labels. MVOcC, medio ventral occipital cortex; FuG, fusiform gyrus; PhG, parahippocampal gyrus; STG, superior temporal gyrus; MTG, middle temporal gyrus; pSTS, posterior superior temporal sulcus; SFG, superior frontal gyrus; CG, cingulate gyrus; SPL, superior parietal lobule.

**Table S6. Cluster statistics and Brainnetome atlas parcellation of ISFC divergence clusters anchored to the dmPFC-rACC seed.**

| Cluster level | | | | | |
|---|---|---|---|---|---|
| **Cluster label** | **Description** | **Size** | **MNI (X,Y,Z) (peak)** | **Fisher's $z$-value (peak)** | **$t$-value (peak)** |
| Cluster-1 | *IFG-OrG-Anterior Insula* | 873 | (-34, 20, -14) | 0.018 | 6.032 |
| Cluster-2 | *Left ventral striatum* | 112 | (-18, 16, -8) | 0.015 | 6.165 |
| Cluster-3 | *IFG-OrG-Anterior Insula* | 330 | (36, 24, 12) | 0.014 | 4.929 |
| Cluster-4 | *IFG-OrG-Anterior Insula* | 1467 | (-32, 48, 32) | 0.018 | 6.866 |
| Cluster-5 | *IFG-OrG-Anterior Insula* | 3 | (-20, 66, -8) | 0.012 | 3.898 |
| Cluster-6 | *Medial prefrontal-cingulate cortex* | 7047 | (-10, 50, 10) | 0.018 | 6.514 |
| Cluster-7 | *IFG-OrG-Anterior Insula* | 16 | (34, 32, 2) | 0.012 | 4.590 |
| Cluster-8 | *IFG-OrG-Anterior Insula* | 16 | (32, 12, 12) | 0.012 | 3.789 |
| Cluster-9 | *Dorsolateral prefrontal cortex* | 664 | (28, 50, 20) | 0.021 | 7.774 |
| Cluster-10 | *A40c, caudal area 40(PFm)* | 13 | (-56, -48, 30) | 0.012 | 4.834 |
| Cluster-11 | *A40c, caudal area 40(PFm)* | 4 | (-64, -50, 32) | 0.011 | 4.573 |

| Voxel level (Cluster-1) | | | | | | |
|---|---|---|---|---|---|---|
| **Atlas ID** | **Name** | **$N_{voxel}$** | **Percent** | **Mean $t$ (range)** | **Anatomical and modified Cyto-architectonic descriptions** | **Gyrus** |
| 33 | A45c_L | 35 | 4.009 | 3.360 (2.861–3.967) | *A45c, caudal area 45* | *IFG* |
| 35 | A45r_L | 6 | 0.687 | 3.405 (3.014–3.738) | *A45r, rostral area 45* | *IFG* |
| 37 | A44op_L | 182 | 20.848 | 3.836 (2.857–5.360) | *A44op, opercular area 44* | *IFG* |
| 39 | A44v_L | 80 | 9.164 | 3.797 (2.883–5.320) | *A44v, ventral area 44* | *IFG* |
| 43 | A12/47o_L | 42 | 4.811 | 4.132 (2.881–6.032) | *A12/47o, orbital area 12/47* | *OrG* |
| 51 | A12/47l_L | 103 | 11.798 | 3.976 (2.862–5.512) | *A12/47l, lateral area 12/47* | *OrG* |
| 61 | A4tl_L | 2 | 0.229 | 3.051 (2.917–3.186) | *A4tl, area 4(tongue and larynx region)* | *PrG* |
| 69 | A38m_L | 1 | 0.115 | 2.869 (2.869–2.869) | *A38m, medial area 38* | *STG* |
| 73 | TE1.0/TE1.2_L | 5 | 0.573 | 3.884 (3.282–4.594) | *TE1.0 and TE1.2* | *STG* |
| 165 | vIa_L | 116 | 13.288 | 4.439 (2.942–5.976) | *vIa, ventral agranular insula* | *INS* |
| 167 | dIa_L | 158 | 18.099 | 3.870 (2.905–5.619) | *dIa, dorsal agranular insula* | *INS* |
| 173 | dId_L | 143 | 16.38 | 3.681 (2.874–5.203) | *dId, dorsal dysgranular insula* | *INS* |

| Voxel level (Cluster-2) | | | | | | |
|---|---|---|---|---|---|---|
| **Atlas ID** | **Name** | **$N_{voxel}$** | **Percent** | **Mean $t$ (range)** | **Anatomical and modified Cyto-architectonic descriptions** | **Gyrus** |
| 49 | A13_L | 11 | 9.821 | 3.806 (3.399–4.780) | *A13, area 13* | *OrG* |
| 219 | vCa_L | 65 | 58.036 | 4.288 (3.334–6.165) | *vCa, ventral caudate* | *BG* |
| 223 | NAC_L | 19 | 16.964 | 4.048 (3.327–5.124) | *NAC, nucleus accumbens* | *BG* |
| 225 | vmPu_L | 17 | 15.179 | 4.041 (3.425–4.356) | *vmPu, ventromedial putamen* | *BG* |

| Voxel level (Cluster-3) | | | | | | |
|---|---|---|---|---|---|---|
| **Atlas ID** | **Name** | **$N_{voxel}$** | **Percent** | **Mean $t$ (range)** | **Anatomical and modified Cyto-architectonic descriptions** | **Gyrus** |
| 34 | A45c_R | 3 | 0.909 | 3.332 (3.250–3.385) | *A45c, caudal area 45* | *IFG* |
| 38 | A44op_R | 163 | 49.394 | 3.863 (3.159–4.929) | *A44op, opercular area 44* | *IFG* |

| | | | | | | |
|---|---|---|---|---|---|---|
| 40 | A44v_R | 7 | 2.121 | 3.385 (3.166–3.594) | *A44v, ventral area 44* | *IFG* |
| 44 | A12/47o_R | 1 | 0.303 | 3.231 (3.231–3.231) | *A12/47o, orbital area 12/47* | *OrG* |
| 46 | A11l_R | 3 | 0.909 | 3.262 (3.163–3.430) | *A11l, lateral area 11* | *OrG* |
| 52 | A12/47l_R | 24 | 7.273 | 3.456 (3.176–3.889) | *A12/47l, lateral area 12/47* | *OrG* |
| 62 | A4tl_R | 5 | 1.515 | 3.636 (3.174–3.931) | *A4tl, area 4(tongue and larynx region)* | *PrG* |
| 74 | TE1.0/TE1.2_R | 34 | 10.303 | 3.510 (3.150–4.070) | *TE1.0 and TE1.2* | *STG* |
| 166 | vIa_R | 30 | 9.091 | 3.632 (3.166–4.645) | *vIa, ventral agranular insula* | *INS* |
| 168 | dIa_R | 50 | 15.152 | 3.625 (3.194–4.609) | *dIa, dorsal agranular insula* | *INS* |
| 174 | dId_R | 10 | 3.03 | 3.523 (3.235–4.026) | *dId, dorsal dysgranular insula* | *INS* |
| **Voxel level (Cluster-4)** | | | | | | |
| **Atlas ID** | **Name** | $N_{voxel}$ | **Percent** | **Mean $t$ (range)** | **Anatomical and modified Cyto-architectonic descriptions** | **Gyrus** |
| 3 | A8dl_L | 16 | 1.091 | 3.164 (2.645–3.848) | *A8dl, dorsolateral area 8* | *SFG* |
| 5 | A9l_L | 74 | 5.044 | 3.740 (2.485–5.924) | *A9l, lateral area 9* | *SFG* |
| 15 | A9/46d_L | 567 | 38.65 | 4.260 (2.509–6.866) | *A9/46d, dorsal area 9/46* | *MFG* |
| 17 | IFJ_L | 19 | 1.295 | 2.864 (2.441–4.168) | *IFJ, inferior frontal junction* | *MFG* |
| 19 | A46_L | 264 | 17.996 | 3.415 (2.435–4.639) | *A46, area 46* | *MFG* |
| 21 | A9/46v_L | 191 | 13.02 | 3.082 (2.436–4.419) | *A9/46v, ventral area 9/46* | *MFG* |
| 23 | A8vl_L | 322 | 21.95 | 4.251 (2.435–6.711) | *A8vl, ventrolateral area 8* | *MFG* |
| 27 | A10l_L | 14 | 0.954 | 2.657 (2.431–3.026) | *A10l, lateral area10* | *MFG* |
| **Voxel level (Cluster-6)** | | | | | | |
| **Atlas ID** | **Name** | $N_{voxel}$ | **Percent** | **Mean $t$ (range)** | **Anatomical and modified Cyto-architectonic descriptions** | **Gyrus** |
| 1 | A8m_L | 377 | 5.35 | 3.450 (2.154–5.945) | *A8m, medial area 8* | *SFG* |
| 2 | A8m_R | 367 | 5.208 | 3.546 (2.158–6.236) | *A8m, medial area 8* | *SFG* |
| 3 | A8dl_L | 133 | 1.887 | 3.816 (2.176–5.772) | *A8dl, dorsolateral area 8* | *SFG* |
| 4 | A8dl_R | 85 | 1.206 | 3.018 (2.164–4.861) | *A8dl, dorsolateral area 8* | *SFG* |
| 5 | A9l_L | 1 | 0.014 | 2.569 (2.569–2.569) | *A9l, lateral area 9* | *SFG* |
| 6 | A9l_R | 15 | 0.213 | 2.742 (2.214–3.657) | *A9l, lateral area 9* | *SFG* |
| 7 | A6dl_L | 232 | 3.292 | 3.705 (2.155–5.548) | *A6dl, dorsolateral area 6* | *SFG* |
| 8 | A6dl_R | 243 | 3.448 | 3.137 (2.157–5.022) | *A6dl, dorsolateral area 6* | *SFG* |
| 9 | A6m_L | 203 | 2.881 | 2.894 (2.150–4.096) | *A6m, medial area 6* | *SFG* |
| 10 | A6m_R | 70 | 0.993 | 2.628 (2.161–3.892) | *A6m, medial area 6* | *SFG* |
| 11 | A9m_L | 282 | 4.002 | 3.606 (2.167–6.389) | *A9m,medial area 9* | *SFG* |
| 12 | A9m_R | 354 | 5.023 | 3.789 (2.153–6.080) | *A9m,medial area 9* | *SFG* |
| 13 | A10m_L | 487 | 6.911 | 3.543 (2.152–6.514) | *A10m, medial area 10* | *SFG* |
| 14 | A10m_R | 420 | 5.96 | 3.171 (2.154–5.411) | *A10m, medial area 10* | *SFG* |
| 25 | A6vl_L | 98 | 1.391 | 2.600 (2.150–3.362) | *A6vl, ventrolateral area 6* | *MFG* |
| 26 | A6vl_R | 2 | 0.028 | 2.408 (2.366–2.450) | *A6vl, ventrolateral area 6* | *MFG* |
| 41 | A14m_L | 47 | 0.667 | 2.708 (2.163–4.115) | *A14m, medial area 14* | *OrG* |
| 42 | A14m_R | 29 | 0.412 | 2.652 (2.157–3.911) | *A14m, medial area 14* | *OrG* |
| 55 | A6cdl_L | 133 | 1.887 | 2.559 (2.152–3.287) | *A6cdl, caudal dorsolateral area 6* | *PrG* |

| | | | | | | |
|---|---|---|---|---|---|---|
| 56 | A6cdl_R | 8 | 0.114 | 2.301 (2.190–2.593) | *A6cdl, caudal dorsolateral area 6* | *PrG* |
| 59 | A4t_L | 6 | 0.085 | 2.442 (2.217–2.691) | *A4t, area 4(trunk region)* | *PrG* |
| 65 | A1/2/3ll_L | 13 | 0.184 | 2.476 (2.240–2.875) | *A1/2/3ll, area1/2/3 (lower limb region)* | *PCL* |
| 66 | A1/2/3ll_R | 53 | 0.752 | 2.567 (2.151–3.931) | *A1/2/3ll, area1/2/3 (lower limb region)* | *PCL* |
| 67 | A4ll_L | 16 | 0.227 | 3.112 (2.150–4.685) | *A4ll, area 4, (lower limb region)* | *PCL* |
| 68 | A4ll_R | 43 | 0.61 | 2.968 (2.190–4.467) | *A4ll, area 4, (lower limb region)* | *PCL* |
| 125 | A7r_L | 7 | 0.099 | 2.381 (2.167–2.678) | *A7r, rostral area 7* | *SPL* |
| 127 | A7c_L | 2 | 0.028 | 2.299 (2.223–2.376) | *A7c, caudal area 7* | *SPL* |
| 131 | A7pc_L | 1 | 0.014 | 2.197 (2.197–2.197) | *A7pc, postcentral area 7* | *SPL* |
| 147 | A7m_L | 157 | 2.228 | 2.915 (2.158–4.087) | *A7m, medial area 7(PEp)* | *Pcun* |
| 148 | A7m_R | 121 | 1.717 | 2.748 (2.151–3.745) | *A7m, medial area 7(PEp)* | *Pcun* |
| 149 | A5m_L | 244 | 3.462 | 2.915 (2.156–4.351) | *A5m, medial area 5(PEm)* | *Pcun* |
| 150 | A5m_R | 219 | 3.108 | 3.123 (2.151–4.315) | *A5m, medial area 5(PEm)* | *Pcun* |
| 153 | A31_L | 32 | 0.454 | 3.045 (2.177–4.824) | *A31, area 31 (Lc1)* | *Pcun* |
| 154 | A31_R | 46 | 0.653 | 2.690 (2.162–3.702) | *A31, area 31 (Lc1)* | *Pcun* |
| 175 | A23d_L | 105 | 1.49 | 3.342 (2.152–5.018) | *A23d, dorsal area 23* | *CG* |
| 176 | A23d_R | 71 | 1.008 | 3.430 (2.150–4.897) | *A23d, dorsal area 23* | *CG* |
| 177 | A24rv_L | 119 | 1.689 | 3.410 (2.151–5.661) | *A24rv, rostroventral area 24* | *CG* |
| 178 | A24rv_R | 107 | 1.518 | 2.904 (2.157–4.624) | *A24rv, rostroventral area 24* | *CG* |
| 179 | A32p_L | 330 | 4.683 | 4.036 (2.210–5.951) | *A32p, pregenual area 32* | *CG* |
| 180 | A32p_R | 200 | 2.838 | 4.556 (2.188–6.233) | *A32p, pregenual area 32* | *CG* |
| 181 | A23v_L | 60 | 0.851 | 3.287 (2.157–4.917) | *A23v, ventral area 23* | *CG* |
| 182 | A23v_R | 34 | 0.482 | 2.905 (2.174–3.879) | *A23v, ventral area 23* | *CG* |
| 183 | A24cd_L | 185 | 2.625 | 3.522 (2.172–5.801) | *A24cd, caudodorsal area 24* | *CG* |
| 184 | A24cd_R | 129 | 1.831 | 3.193 (2.172–4.929) | *A24cd, caudodorsal area 24* | *CG* |
| 185 | A23c_L | 341 | 4.839 | 3.352 (2.156–5.029) | *A23c, caudal area 23* | *CG* |
| 186 | A23c_R | 307 | 4.356 | 3.498 (2.166–5.147) | *A23c, caudal area 23* | *CG* |
| 187 | A32sg_L | 248 | 3.519 | 3.411 (2.156–5.348) | *A32sg, subgenual area 32* | *CG* |
| 188 | A32sg_R | 265 | 3.76 | 3.664 (2.160–5.629) | *A32sg, subgenual area 32* | *CG* |

**Voxel level (Cluster-9)**

| Atlas ID | Name | $N_{voxel}$ | Percent | Mean $t$ (range) | Anatomical and modified Cyto-architectonic descriptions | Gyrus |
|---|---|---|---|---|---|---|
| 4 | A8dl_R | 12 | 1.807 | 3.550 (3.199–4.149) | *A8dl, dorsolateral area 8* | *SFG* |
| 6 | A9l_R | 35 | 5.271 | 3.465 (3.156–4.603) | *A9l, lateral area 9* | *SFG* |
| 16 | A9/46d_R | 337 | 50.753 | 4.203 (3.145–6.150) | *A9/46d, dorsal area 9/46* | *MFG* |
| 20 | A46_R | 279 | 42.018 | 4.842 (3.145–7.774) | *A46, area 46* | *MFG* |
| 22 | A9/46v_R | 1 | 0.151 | 3.685 (3.685–3.685) | *A9/46v, ventral area 9/46* | *MFG* |

The upper section (cluster level) summarizes all voxel clusters identified in the whole-brain ISFC divergence analysis using the dmPFC-rACC as a seed region. For each cluster, the table lists the cluster label, anatomical description, size (voxels), MNI peak coordinates (x, y, z; mm), peak Fisher's z-transformed ISFC difference, and peak voxel-wise $t$-statistic. All clusters survived TFCE-corrected $P < 0.05$ (10,000 permutations), reflecting significantly stronger within-group than between-group stimulus-driven coupling with the dmPFC-rACC. The lower sections (voxel level) detail the Brainnetome atlas parcellation for each cluster exceeding 20 voxels; clusters with 20 or fewer voxels are listed in the cluster-level summary only. For each parcel, the Atlas ID and anatomical name follow the Brainnetome atlas nomenclature. $N_{voxel}$ gives the number of cluster voxels falling within each parcel; % of cluster is the corresponding proportion of the cluster total. Mean $t$ (range) reports the mean voxel-wise t-statistic across all voxels in that parcel, with the minimum and maximum values shown in parentheses. Gyrus and Lobe indicate macro-anatomical labels. MNI, Montreal Neurological Institute; ISFC, inter-subject functional connectivity; TFCE, threshold-free cluster enhancement; BG, basal ganglia; CG, cingulate gyrus; IFG, inferior frontal gyrus; INS, insular gyrus; IPL, inferior parietal lobule; MFG, middle frontal gyrus; OrG, orbital gyrus; PCL, paracentral lobule; Pcun, precuneus; PrG, precentral gyrus; SFG, superior frontal Gyrus; SPL, superior parietal lobule; STG, superior temporal gyrus.

**Table S7. Cluster statistics and Brainnetome atlas parcellation of ISFC divergence clusters anchored to the left dlPFC seed.**

| Cluster level | | | | | |
|---|---|---|---|---|---|
| **Cluster label** | **Description** | **Size** | **MNI (X,Y,Z) (peak)** | **Fisher's $z$-value (peak)** | **$t$-value (peak)** |
| Cluster-1 | *IFG-OrG-Anterior Insula Complex & Striatum* | 1909 | (-36, 16, -12) | 0.021 | 6.717 |
| Cluster-2 | *IFG-OrG-Anterior Insula Complex* | 76 | (26, 20, -12) | 0.011 | 3.745 |
| Cluster-3 | *Medial prefrontal-cingulate cortex & Dorsolateral prefrontal cortex* | 8824 | (-28, 54, 26) | 0.023 | 6.923 |
| Cluster-4 | *Striatum* | 210 | (18, 26, 2) | 0.017 | 5.695 |
| Cluster-5 | *IFG-OrG-Anterior Insula Complex* | 15 | (-2, 10, -8) | 0.011 | 4.372 |
| Cluster-6 | *IFG-OrG-Anterior Insula Complex* | 361 | (52, 8, -2) | 0.013 | 4.384 |
| Cluster-7 | *Caudal ventrolateral area 6* | 29 | (-58, 8, 16) | 0.008 | 3.563 |
| Cluster-8 | *Dorsolateral prefrontal cortex* | 736 | (34, 54, 24) | 0.018 | 6.581 |
| Cluster-9 | *Inferior parietal lobule* | 553 | (-62, -54, 30) | 0.018 | 6.476 |
| Cluster-10 | *Caudal ventrolateral area 6* | 56 | (-54, 6, 36) | 0.012 | 4.372 |
| Cluster-11 | *Inferior parietal lobule* | 107 | (56, -54, 46) | 0.013 | 6.090 |
| Cluster-12 | *Medial prefrontal-cingulate cortex* | 23 | (-8, -52, 58) | 0.013 | 5.906 |

| Voxel level (Cluster-1) | | | | | | |
|---|---|---|---|---|---|---|
| **Atlas ID** | **Name** | **$N_{voxel}$** | **Percent** | **Mean $t$ (range)** | **Anatomical and modified Cyto-architectonic descriptions** | **Gyrus** |
| 33 | A45c_L | 73 | 3.824 | 3.386 (2.433–4.999) | *A45c, caudal area 45* | *IFG* |
| 35 | A45r_L | 28 | 1.467 | 3.114 (2.414–4.099) | *A45r, rostral area 45* | *IFG* |
| 37 | A44op_L | 216 | 11.315 | 3.480 (2.419–5.477) | *A44op, opercular area 44* | *IFG* |
| 39 | A44v_L | 132 | 6.915 | 3.186 (2.413–4.742) | *A44v, ventral area 44* | *IFG* |
| 43 | A12/47o_L | 117 | 6.129 | 4.241 (2.496–6.569) | *A12/47o, orbital area 12/47* | *OrG* |
| 45 | A11l_L | 142 | 7.438 | 3.654 (2.467–5.143) | *A11l, lateral area 11* | *OrG* |
| 49 | A13_L | 93 | 4.872 | 3.046 (2.411–4.266) | *A13, area 13* | *OrG* |
| 51 | A12/47l_L | 173 | 9.062 | 4.205 (2.454–6.425) | *A12/47l, lateral area 12/47* | *OrG* |
| 61 | A4tl_L | 1 | 0.052 | 3.249 (3.249–3.249) | *A4tl, area 4(tongue and larynx region)* | *PrG* |
| 63 | A6cvl_L | 2 | 0.105 | 2.488 (2.471–2.505) | *A6cvl, caudal ventrolateral area 6* | *PrG* |
| 69 | A38m_L | 6 | 0.314 | 3.115 (2.791–3.313) | *A38m, medial area 38* | *STG* |
| 73 | TE1.0/TE1.2_L | 4 | 0.21 | 2.972 (2.415–3.673) | *TE1.0 and TE1.2* | *STG* |
| 77 | A38l_L | 18 | 0.943 | 3.231 (2.472–4.043) | *A38l, lateral area 38* | *STG* |
| 165 | vIa_L | 182 | 9.534 | 4.408 (2.426–6.717) | *vIa, ventral agranular insula* | *INS* |
| 167 | dIa_L | 162 | 8.486 | 4.187 (2.422–6.436) | *dIa, dorsal agranular insula* | *INS* |
| 169 | vId/vIg_L | 5 | 0.262 | 2.875 (2.452–3.247) | *vId/vIg, ventral dysgranular and granular insula* | *INS* |
| 173 | dId_L | 186 | 9.743 | 4.009 (2.412–5.742) | *dId, dorsal dysgranular insula* | *INS* |
| 219 | vCa_L | 212 | 11.105 | 3.928 (2.414–6.409) | *vCa, ventral caudate* | *BG* |
| 223 | NAC_L | 75 | 3.929 | 3.198 (2.444–4.663) | *NAC, nucleus accumbens* | *BG* |
| 225 | vmPu_L | 34 | 1.781 | 3.470 (2.497–5.146) | *vmPu, ventromedial putamen* | *BG* |

| | | | | | | |
|---|---|---|---|---|---|---|
| 227 | dCa_L | 48 | 2.514 | 3.164 (2.412–4.121) | *dCa, dorsal caudate* | *BG* |

**Voxel level (Cluster-2)**

| **Atlas ID** | **Name** | $N_{voxel}$ | **Percent** | **Mean $t$ (range)** | **Anatomical and modified Cyto-architectonic descriptions** | **Gyrus** |
|---|---|---|---|---|---|---|
| 44 | A12/47o_R | 4 | 5.263 | 3.157 (2.931–3.389) | *A12/47o, orbital area 12/47* | *OrG* |
| 46 | A11l_R | 20 | 26.316 | 3.243 (2.941–3.745) | *A11l, lateral area 11* | *OrG* |
| 52 | A12/47l_R | 20 | 26.316 | 3.190 (2.950–3.631) | *A12/47l, lateral area 12/47* | *OrG* |
| 166 | vIa_R | 32 | 42.105 | 3.182 (2.934–3.604) | *vIa, ventral agranular insula* | *INS* |

**Voxel level (Cluster-3)**

| **Atlas ID** | **Name** | $N_{voxel}$ | **Percent** | **Mean $t$ (range)** | **Anatomical and modified Cyto-architectonic descriptions** | **Gyrus** |
|---|---|---|---|---|---|---|
| 1 | A8m_L | 527 | 5.972 | 3.726 (2.098–6.062) | *A8m, medial area 8* | *SFG* |
| 2 | A8m_R | 550 | 6.233 | 3.257 (2.109–5.027) | *A8m, medial area 8* | *SFG* |
| 3 | A8dl_L | 190 | 2.153 | 4.051 (2.101–6.776) | *A8dl, dorsolateral area 8* | *SFG* |
| 4 | A8dl_R | 81 | 0.918 | 3.140 (2.112–4.581) | *A8dl, dorsolateral area 8* | *SFG* |
| 5 | A9l_L | 190 | 2.153 | 3.572 (2.090–6.539) | *A9l, lateral area 9* | *SFG* |
| 6 | A9l_R | 14 | 0.159 | 2.774 (2.165–3.500) | *A9l, lateral area 9* | *SFG* |
| 7 | A6dl_L | 211 | 2.391 | 3.535 (2.094–5.693) | *A6dl, dorsolateral area 6* | *SFG* |
| 8 | A6dl_R | 85 | 0.963 | 3.276 (2.098–4.852) | *A6dl, dorsolateral area 6* | *SFG* |
| 9 | A6m_L | 158 | 1.791 | 3.105 (2.103–5.135) | *A6m, medial area 6* | *SFG* |
| 10 | A6m_R | 109 | 1.235 | 2.636 (2.092–3.700) | *A6m, medial area 6* | *SFG* |
| 11 | A9m_L | 307 | 3.479 | 3.689 (2.093–6.693) | *A9m,medial area 9* | *SFG* |
| 12 | A9m_R | 408 | 4.624 | 3.740 (2.101–5.782) | *A9m,medial area 9* | *SFG* |
| 13 | A10m_L | 651 | 7.378 | 3.348 (2.094–5.759) | *A10m, medial area 10* | *SFG* |
| 14 | A10m_R | 474 | 5.372 | 3.306 (2.114–5.337) | *A10m, medial area 10* | *SFG* |
| 15 | A9/46d_L | 534 | 6.052 | 4.293 (2.094–6.923) | *A9/46d, dorsal area 9/46* | *MFG* |
| 17 | IFJ_L | 25 | 0.283 | 2.801 (2.110–4.021) | *IFJ, inferior frontal junction* | *MFG* |
| 19 | A46_L | 450 | 5.1 | 3.964 (2.107–6.810) | *A46, area 46* | *MFG* |
| 21 | A9/46v_L | 201 | 2.278 | 2.890 (2.097–4.838) | *A9/46v, ventral area 9/46* | *MFG* |
| 23 | A8vl_L | 324 | 3.672 | 3.656 (2.094–6.027) | *A8vl, ventrolateral area 8* | *MFG* |
| 25 | A6vl_L | 82 | 0.929 | 2.724 (2.126–3.561) | *A6vl, ventrolateral area 6* | *MFG* |
| 27 | A10l_L | 365 | 4.136 | 3.571 (2.109–5.827) | *A10l, lateral area10* | *MFG* |
| 35 | A45r_L | 16 | 0.181 | 2.431 (2.115–2.820) | *A45r, rostral area 45* | *IFG* |
| 41 | A14m_L | 73 | 0.827 | 2.554 (2.093–4.442) | *A14m, medial area 14* | *OrG* |
| 42 | A14m_R | 92 | 1.043 | 2.975 (2.112–4.024) | *A14m, medial area 14* | *OrG* |
| 43 | A12/47o_L | 61 | 0.691 | 3.131 (2.096–5.916) | *A12/47o, orbital area 12/47* | *OrG* |
| 45 | A11l_L | 45 | 0.51 | 2.688 (2.115–4.287) | *A11l, lateral area 11* | *OrG* |
| 47 | A11m_L | 22 | 0.249 | 2.876 (2.097–4.181) | *A11m, medial area 11* | *OrG* |
| 49 | A13_L | 5 | 0.057 | 2.470 (2.163–2.677) | *A13, area 13* | *OrG* |
| 51 | A12/47l_L | 19 | 0.215 | 3.139 (2.233–5.203) | *A12/47l, lateral area 12/47* | *OrG* |
| 55 | A6cdl_L | 19 | 0.215 | 2.624 (2.112–3.270) | *A6cdl, caudal dorsolateral area 6* | *PrG* |
| 66 | A1/2/3ll_R | 46 | 0.521 | 2.634 (2.093–4.251) | *A1/2/3ll, area1/2/3 (lower limb region)* | *PCL* |

| | | | | | | |
|---|---|---|---|---|---|---|
| 67 | A4ll_L | 1 | 0.011 | 3.388 (3.388–3.388) | *A4ll, area 4, (lower limb region)* | *PCL* |
| 68 | A4ll_R | 19 | 0.215 | 2.671 (2.135–4.088) | *A4ll, area 4, (lower limb region)* | *PCL* |
| 149 | A5m_L | 13 | 0.147 | 2.541 (2.160–2.886) | *A5m, medial area 5(PEm)* | *Pcun* |
| 150 | A5m_R | 154 | 1.745 | 2.891 (2.102–4.852) | *A5m, medial area 5(PEm)* | *Pcun* |
| 153 | A31_L | 12 | 0.136 | 2.478 (2.133–3.038) | *A31, area 31 (Lc1)* | *Pcun* |
| 154 | A31_R | 16 | 0.181 | 2.596 (2.126–3.011) | *A31, area 31 (Lc1)* | *Pcun* |
| 175 | A23d_L | 67 | 0.759 | 3.200 (2.282–4.155) | *A23d, dorsal area 23* | *CG* |
| 176 | A23d_R | 71 | 0.805 | 3.220 (2.105–4.267) | *A23d, dorsal area 23* | *CG* |
| 177 | A24rv_L | 64 | 0.725 | 3.354 (2.119–5.195) | *A24rv, rostroventral area 24* | *CG* |
| 178 | A24rv_R | 107 | 1.213 | 3.141 (2.091–4.897) | *A24rv, rostroventral area 24* | *CG* |
| 179 | A32p_L | 326 | 3.694 | 4.127 (2.254–6.804) | *A32p, pregenual area 32* | *CG* |
| 180 | A32p_R | 200 | 2.267 | 4.682 (2.414–6.061) | *A32p, pregenual area 32* | *CG* |
| 181 | A23v_L | 6 | 0.068 | 2.638 (2.130–3.218) | *A23v, ventral area 23* | *CG* |
| 182 | A23v_R | 2 | 0.023 | 2.521 (2.436–2.605) | *A23v, ventral area 23* | *CG* |
| 183 | A24cd_L | 169 | 1.915 | 3.197 (2.100–5.421) | *A24cd, caudodorsal area 24* | *CG* |
| 184 | A24cd_R | 147 | 1.666 | 3.246 (2.117–4.989) | *A24cd, caudodorsal area 24* | *CG* |
| 185 | A23c_L | 222 | 2.516 | 2.945 (2.095–4.614) | *A23c, caudal area 23* | *CG* |
| 186 | A23c_R | 220 | 2.493 | 3.053 (2.090–4.540) | *A23c, caudal area 23* | *CG* |
| 187 | A32sg_L | 375 | 4.25 | 3.686 (2.101–6.101) | *A32sg, subgenual area 32* | *CG* |
| 188 | A32sg_R | 299 | 3.388 | 4.254 (2.133–6.745) | *A32sg, subgenual area 32* | *CG* |
| **Voxel level (Cluster-4)** | | | | | | |
| **Atlas ID** | **Name** | $N_{voxel}$ | **Percent** | **Mean $t$ (range)** | **Anatomical and modified Cyto-architectonic descriptions** | **Gyrus** |
| 220 | vCa_R | 78 | 37.143 | 3.480 (2.790–5.257) | *vCa, ventral caudate* | *BG* |
| 224 | NAC_R | 26 | 12.381 | 3.384 (2.746–4.327) | *NAC, nucleus accumbens* | *BG* |
| 226 | vmPu_R | 6 | 2.857 | 2.919 (2.758–3.208) | *vmPu, ventromedial putamen* | *BG* |
| 228 | dCa_R | 100 | 47.619 | 3.703 (2.733–5.695) | *dCa, dorsal caudate* | *BG* |
| **Voxel level (Cluster-5)** | | | | | | |
| **Atlas ID** | **Name** | $N_{voxel}$ | **Percent** | **Mean $t$ (range)** | **Anatomical and modified Cyto-architectonic descriptions** | **Gyrus** |
| 49 | A13_L | 14 | 93.333 | 3.774 (3.285–4.372) | *A13, area 13* | *OrG* |
| 50 | A13_R | 1 | 6.667 | 3.431 (3.431–3.431) | *A13, area 13* | *OrG* |
| **Voxel level (Cluster-6)** | | | | | | |
| **Atlas ID** | **Name** | $N_{voxel}$ | **Percent** | **Mean $t$ (range)** | **Anatomical and modified Cyto-architectonic descriptions** | **Gyrus** |
| 34 | A45c_R | 56 | 15.512 | 3.282 (2.709–4.248) | *A45c, caudal area 45* | *IFG* |
| 36 | A45r_R | 16 | 4.432 | 3.232 (2.718–4.114) | *A45r, rostral area 45* | *IFG* |
| 38 | A44op_R | 159 | 44.044 | 3.307 (2.668–4.325) | *A44op, opercular area 44* | *IFG* |
| 40 | A44v_R | 16 | 4.432 | 3.303 (2.801–3.813) | *A44v, ventral area 44* | *IFG* |
| 52 | A12/47l_R | 5 | 1.385 | 3.035 (2.684–3.389) | *A12/47l, lateral area 12/47* | *OrG* |
| 62 | A4tl_R | 12 | 3.324 | 3.417 (2.749–4.327) | *A4tl, area 4(tongue and larynx region)* | *PrG* |
| 74 | TE1.0/TE1.2_R | 56 | 15.512 | 3.445 (2.674–4.384) | *TE1.0 and TE1.2* | *STG* |

| Atlas ID | Name | $N_{voxel}$ | Percent | Mean $t$ (range) | Anatomical and modified Cyto-architectonic descriptions | Gyrus |
|---|---|---|---|---|---|---|
| 168 | dIa_R | 26 | 7.202 | 3.323 (2.711–4.189) | *dIa, dorsal agranular insula* | *INS* |
| 174 | dId_R | 15 | 4.155 | 3.412 (2.810–3.983) | *dId, dorsal dysgranular insula* | *INS* |
| **Voxel level (Cluster-7)** | | | | | | |
| **Atlas ID** | **Name** | $N_{voxel}$ | **Percent** | **Mean $t$ (range)** | **Anatomical and modified Cyto-architectonic descriptions** | **Gyrus** |
| 53 | A4hf_L | 1 | 3.448 | 2.678 (2.678–2.678) | *A4hf, area 4(head and face region)* | *PrG* |
| 63 | A6cvl_L | 28 | 96.552 | 2.959 (2.622–3.563) | *A6cvl, caudal ventrolateral area 6* | *PrG* |
| **Voxel level (Cluster-8)** | | | | | | |
| **Atlas ID** | **Name** | $N_{voxel}$ | **Percent** | **Mean $t$ (range)** | **Anatomical and modified Cyto-architectonic descriptions** | **Gyrus** |
| 4 | A8dl_R | 8 | 1.087 | 3.750 (3.196–4.620) | *A8dl, dorsolateral area 8* | *SFG* |
| 6 | A9l_R | 29 | 3.94 | 3.763 (3.212–5.171) | *A9l, lateral area 9* | *SFG* |
| 16 | A9/46d_R | 347 | 47.147 | 4.295 (3.194–6.252) | *A9/46d, dorsal area 9/46* | *MFG* |
| 20 | A46_R | 328 | 44.565 | 4.667 (3.193–6.581) | *A46, area 46* | *MFG* |
| 22 | A9/46v_R | 23 | 3.125 | 4.110 (3.303–5.574) | *A9/46v, ventral area 9/46* | *MFG* |
| 24 | A8vl_R | 1 | 0.136 | 3.338 (3.338–3.338) | *A8vl, ventrolateral area 8* | *MFG* |
| **Voxel level (Cluster-9)** | | | | | | |
| **Atlas ID** | **Name** | $N_{voxel}$ | **Percent** | **Mean $t$ (range)** | **Anatomical and modified Cyto-architectonic descriptions** | **Gyrus** |
| 137 | A39rd_L | 7 | 1.266 | 3.578 (3.297–3.953) | *A39rd, rostrodorsal area 39(Hip3)* | *IPL* |
| 139 | A40rd_L | 18 | 3.255 | 3.713 (3.266–4.728) | *A40rd, rostrodorsal area 40(PFt)* | *IPL* |
| 141 | A40c_L | 510 | 92.224 | 4.324 (3.262–6.476) | *A40c, caudal area 40(PFm)* | *IPL* |
| 143 | A39rv_L | 7 | 1.266 | 3.951 (3.365–4.680) | *A39rv, rostroventral area 39(PGa)* | *IPL* |
| 145 | A40rv_L | 11 | 1.989 | 3.781 (3.284–4.399) | *A40rv, rostroventral area 40(PFop)* | *IPL* |
| **Voxel level (Cluster-10)** | | | | | | |
| **Atlas ID** | **Name** | $N_{voxel}$ | **Percent** | **Mean $t$ (range)** | **Anatomical and modified Cyto-architectonic descriptions** | **Gyrus** |
| 53 | A4hf_L | 2 | 3.571 | 2.732 (2.712–2.751) | *A4hf, area 4(head and face region)* | *PrG* |
| 63 | A6cvl_L | 54 | 96.429 | 3.311 (2.543–4.372) | *A6cvl, caudal ventrolateral area 6* | *PrG* |
| **Voxel level (Cluster-11)** | | | | | | |
| **Atlas ID** | **Name** | $N_{voxel}$ | **Percent** | **Mean $t$ (range)** | **Anatomical and modified Cyto-architectonic descriptions** | **Gyrus** |
| 142 | A40c_R | 57 | 53.271 | 5.159 (4.375–6.090) | *A40c, caudal area 40(PFm)* | *IPL* |
| 144 | A39rv_R | 50 | 46.729 | 4.918 (4.379–5.609) | *A39rv, rostroventral area 39(PGa)* | *IPL* |
| **Voxel level (Cluster-12)** | | | | | | |
| **Atlas ID** | **Name** | $N_{voxel}$ | **Percent** | **Mean $t$ (range)** | **Anatomical and modified Cyto-architectonic descriptions** | **Gyrus** |
| 149 | A5m_L | 23 | 100 | 4.341 (3.664–5.906) | A5m, medial area 5(PEm) | Pcun |

The upper section (cluster level) summarizes all voxel clusters identified in the whole-brain ISFC divergence analysis using the left dlPFC as a seed region. For each cluster, the table lists the cluster label, anatomical description, size (voxels), MNI peak coordinates (x, y, z; mm), peak Fisher's z-transformed ISFC difference, and peak voxel-wise $t$-statistic. All clusters survived TFCE-corrected $P < 0.05$ (10,000 permutations), reflecting significantly stronger within-group than between-group stimulus-driven coupling with the left dlPFC. For each parcel, the Atlas ID and anatomical name follow the Brainnetome atlas nomenclature. $N_{voxel}$ gives the number of cluster voxels falling within each parcel; % of cluster is the corresponding proportion of the cluster total. Mean $t$ (range) reports the mean voxel-wise t-statistic across all voxels in that parcel, with the minimum and maximum values shown in parentheses. Gyrus and Lobe indicate macro-anatomical labels. MNI, Montreal Neurological Institute; ISFC, inter-subject functional connectivity; TFCE, threshold-free cluster enhancement; BG, basal ganglia; CG, cingulate gyrus; IFG, inferior frontal gyrus; INS, insular gyrus; IPL, inferior parietal lobule; MFG, middle frontal gyrus; OrG, orbital gyrus; PCL, paracentral lobule; Pcun, precuneus; PrG, precentral gyrus; SFG, superior frontal gyrus; SPL, superior parietal lobule; STG, superior temporal gyrus.

**Table S8. Summary of phenotypic measures used in this study**

| Domain | Abbreviations of measures | Item |
|---|---|---|
| Demographics | Basic_Demos | EID, Sex |
| Scan information | MRI_Track | Age_at_Scan, Scan_Location |
| Medical status | DrugScreen | AMP500, BAR300, BUP10, BZO300, COC150, MAMP500, MDMA500, MTD300, OPI300, OXY100, PCP25, PPX300, TCA1000, THC50 |
| Clinical diagnosis | Diagnosis_ClinicianConsensus | DX_01~10, DX_01_Time~DX_10_Time |
| Social, emotional, and behavioral function | MFQ_SR | MFQ_SR_Total |

### References

1. Kriegeskorte, N., M. Mur, and P.A. Bandettini, *Representational similarity analysis-connecting the branches of systems neuroscience.* Frontiers in systems neuroscience, 2008. **2**: p. 249.
2. Yan, C.-G., et al., *DPABI: data processing & analysis for (resting-state) brain imaging.* Neuroinformatics, 2016. **14**(3): p. 339-351.
3. Winkler, A.M., et al., *Faster permutation inference in brain imaging.* Neuroimage, 2016. **141**: p. 502-516.
4. Fan, L., et al., *The human brainnetome atlas: a new brain atlas based on connectional architecture.* Cerebral cortex, 2016. **26**(8): p. 3508-3526.
